# Systematic Dissection of Sequence Features Affecting the Binding Specificity of a Pioneer Factor Reveals Binding Synergy Between FOXA1 and AP-1

**DOI:** 10.1101/2023.11.08.566246

**Authors:** Cheng Xu, Holly Kleinschmidt, Jianyu Yang, Erik Leith, Jenna Johnson, Song Tan, Shaun Mahony, Lu Bai

## Abstract

Despite the unique ability of pioneer transcription factors (PFs) to target nucleosomal sites in closed chromatin, they only bind a small fraction of their genomic motifs. The underlying mechanism of this selectivity is not well understood. Here, we design a high-throughput assay called ChIP-ISO to systematically dissect sequence features affecting the binding specificity of a classic PF, FOXA1. Combining ChIP-ISO with *in vitro* and neural network analyses, we find that 1) FOXA1 binding is strongly affected by co-binding TFs AP-1 and CEBPB, 2) FOXA1 and AP-1 show binding cooperativity *in vitro*, 3) FOXA1’s binding is determined more by local sequences than chromatin context, including eu-/heterochromatin, and 4) AP-1 is partially responsible for differential binding of FOXA1 in different cell types. Our study presents a framework for elucidating genetic rules underlying PF binding specificity and reveals a mechanism for context-specific regulation of its binding.

## Main

Sequence-specific transcription factors (TFs) are major regulators of gene expression. Characterization of the location and strength of TF binding in the genome is therefore a critical step in understanding gene regulation. TF binding sites are typically identified using the weight matrices of their binding motifs. In higher eukaryotes, however, this method has weak predictive power for actual TF binding events. Many TFs bind <1% of their motifs across the genome, and their binding patterns can change in a cell-type-specific manner^1–3^. Multiple features beyond the core sequence motif have been proposed to contribute to this phenomenon, including DNA shape^4,5^, cooperative binding with other TFs^6–8^, DNA methylation^9,10^, nucleosome occupancy and chromatin accessibility^11,12^, histone modifications^13,14^, 3D genome contacts^15^, and variations in local TF concentrations^16,17^. Among these potential factors, nucleosomes have a major inhibitory effect on the binding of many TFs, and chromatin accessibility is therefore considered to be the key determinant of TF binding^12,18–20^.

A subset of TFs known as “pioneer factors” (PFs) can stably associate with nucleosomal templates by recognizing partial sequence motifs and/or interacting with histones^14,21–24^. Inside cells, PFs can overcome the nucleosomal barrier by targeting nucleosome-embedded motifs and generating accessible chromatin, which enables the binding of other TFs and triggers transcriptional activation^25–28^. Given their ability to open chromatin *in vivo* and bind nucleosomal DNA *in vitro*, PFs should be able to access most, if not all, consensus motifs in the genome. This is indeed the case for PFs in budding yeast^29^ (**Extended Data Fig. 1a**). Surprisingly, like canonical TFs, PFs in higher eukaryotes also show highly selective and cell-type specific binding. For example, FOXA1 is a classic PF capable of binding and opening highly compacted chromatin^30–33^, but it only occupies 3.7% of its potential motifs in MCF-7 cells, and less than half of these binding events are shared with LNCaP cells^34^. Our analysis found that only 10-20% consensus motifs are bound by FOXA1 in MCF-7 and A549 cells (**Extended Data Fig. 1b**). The molecular mechanism underlying such binding selectivity is not well understood.

TF binding is usually studied in the context of the native genome, where each binding event can be affected by multiple variables, and individual effects are therefore hard to dissect. Here, we sought to overcome this limitation by developing a new method named “Chromatin Immunoprecipitation with Integrated Synthetic Oligonucleotides (ChIP-ISO)” and applied this method to study FOXA1 binding in human A549 lung cancer cells. In this method, we engineered specific genetic features into synthetic sequences, integrated them into a fixed genomic locus, and measured FOXA1 binding in this highly controlled genetic and epigenetic context. In combination with *in vitro* and neural network analyses, our work reveals key determinants of PF binding, which has implications on PF function through development and differentiation.

## Results

### ChIP-ISO assay allows highly parallel measurements of FOXA1 binding to thousands of integrated synthetic sequences

FOXA1 is expressed in A549 cells, where it plays important physiological roles^35,36^. To study FOXA1 binding specificity in this cell line, we performed the ChIP-ISO procedure as shown in **Fig. 1a**. Briefly, a synthetic oligo library (length: 193bp, complexity: 3,203) was inserted into a truncated *CCND1 enhancer* (*CCND1e*)^37^ where all endogenous FOXA1 motifs are deleted. These sequences contain variations in genetic features that potentially affect FOXA1 binding (**Table S1**). The resulting plasmid library was integrated into the *AAVS1* locus in the human genome through CRISPR-Cas9, and FOXA1 binding to these sequences was measured by ChIP followed by amplicon sequencing (**Extended Data Fig. 2**; **Methods**). The same amplicon sequencing was performed on purified genomic sequences as the input control. Since the synthetic sequences associated with FOXA1 are enriched by ChIP, for each library sequence, the ratio of the read count in the ChIP sample divided by that in the input sample was used as a measure of FOXA1 binding strength (referred to as the “ChIP-ISO signal” below) (**Methods**).

**Fig 1:**
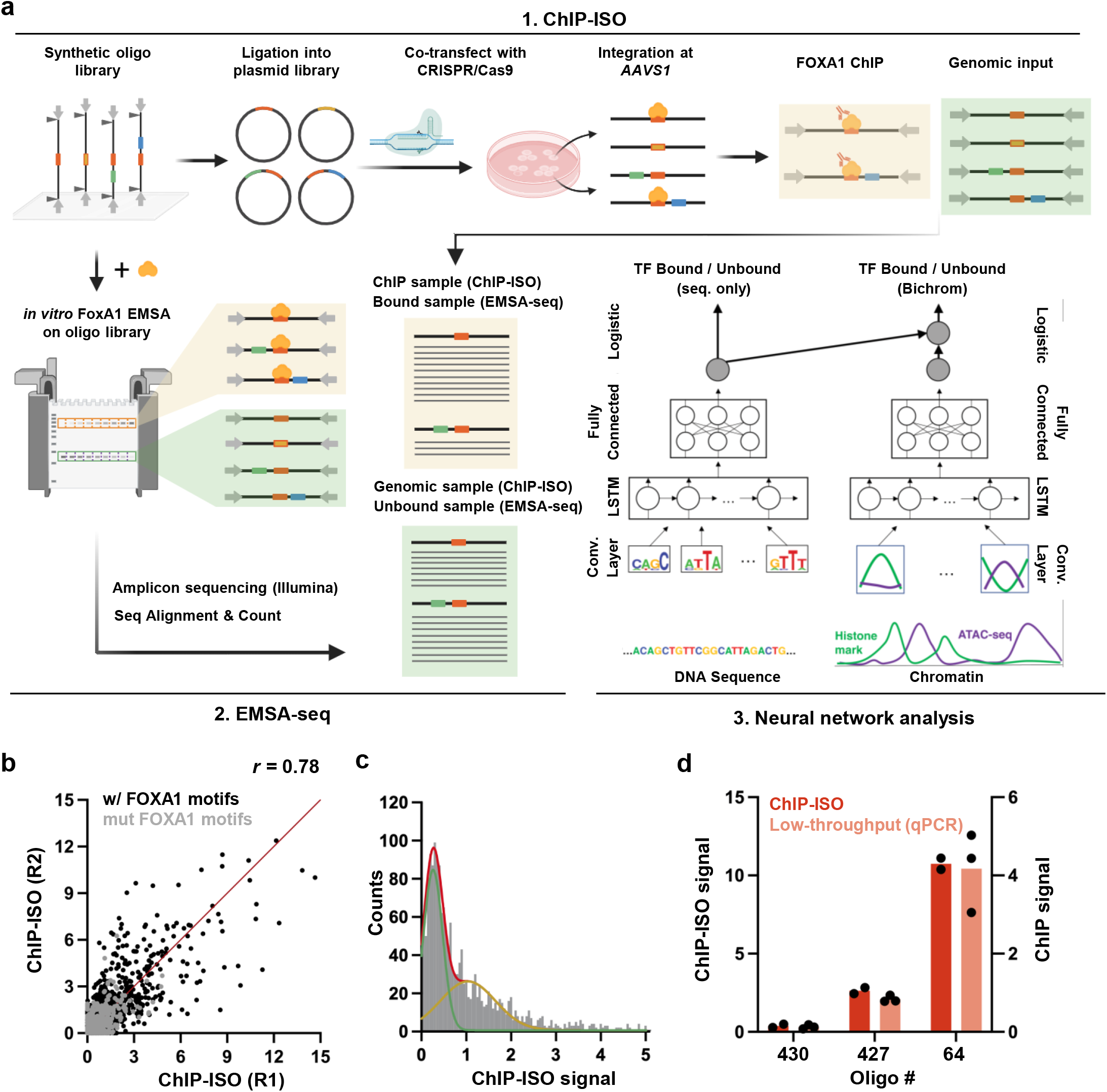
ChIP-ISO assay allows highly parallel measurements of FOXA1 binding to thousands of integrated synthetic sequences. a) Workflows for ChIP-ISO (1), EMSA-seq (2), and neural network analysis (3). b) Reproducibility of FOXA1 ChIP-ISO signal across two biological replicates. Black / gray dots represent sequences containing WT / mutated FOXA1 motifs. r: Pearson correlation. c) Histogram of the FOXA1 ChIP-ISO signals with the entire ISO library, fit by two Gaussian peaks (green and yellow: low and high peaks, respectively; red: superposition of the two). d) Comparison of FOXA1 ChIP-ISO signals with low-throughput ChIP-qPCR signals from three biological replicates over three individual ISO library sequences.

We performed a few tests to evaluate the ChIP-ISO method. Two biological replicates agree well with an overall correlation coefficient of 0.76 (**Fig. 1b**). The ChIP-ISO signals follow a bimodal distribution, with 56.3% of the sequences in the lower peak, representing no or low-level FOXA1 binding (**Fig. 1c**). As expected, sequences with mutated FOXA1 motifs show lower binding (**Fig. 1b**). Furthermore, we constructed three cell lines each containing a single library sequence integrated into the *AAVS1* site, and measured FOXA1 binding by ChIP followed by quantitative PCR (ChIP-qPCR). These low-throughput measurements agree well with the high-throughput results (**Fig. 1d**). We therefore conclude that the ChIP-ISO method can accurately and efficiently measure FOXA1 binding to integrated synthetic sequences.

### Co-binding of AP-1 strongly enhances FOXA1 binding to the *CCND1 enhancer*

The endogenous *CCND1e* in A549 cells is bound by FOXA1 and accompanied by high chromatin accessibility and H3K27ac signals^37^ (**Extended Data Fig. 3a**). It contains three FOXA1 motifs, as well as conserved binding sites of eight other TFs (**Fig. 2a**, **Extended Data Fig. 3b**). The first set of the library includes *CCND1e* mutants. **Fig. 2b** shows the ChIP-ISO measurement on sequences with scanning mutations, where a 10bp window is sequentially scrambled with a 3bp step size. The most prominent drop in FOXA1 occupancy is observed when a region near the third FOXA1 motif is scrambled, indicating that this region contains key elements that recruit FOXA1.

**Fig 2:**
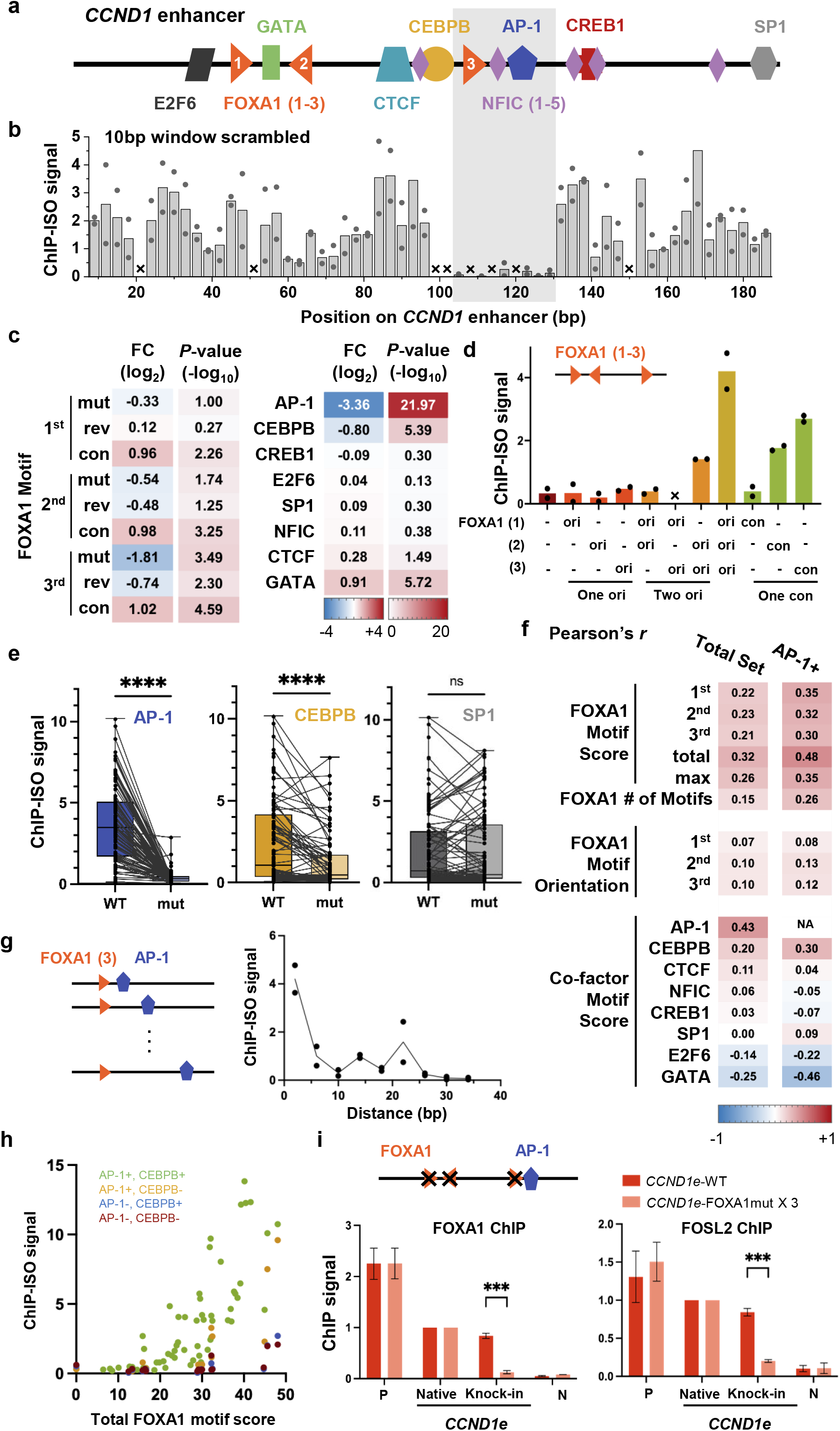
Co-binding of AP-1 strongly enhances FOXA1 binding to the CCND1e enhancer. a) Map of the 193bp portion of the CCND1e explored in this study (chr11:69,654,913-69,655,105). Three FOXA1 motifs (orientations depicted by arrow directions) and motifs of potential co-binding TFs are labeled. b) ChIP-ISO signals over CCND1e variants containing scrambled sequences in a 10bp moving window (step size: 3bp). Bar: averaged ChIP-ISO signal; Dot: data from individual replica; X: missing data. Gray box highlights an area where the scrambles lead to particularly low FOXA1 ChIP-ISO signals, indicating that these sequences are critical for FOXA1 binding. c) The effect on FOXA1 binding by manipulating individual FOXA1 motifs (left) or mutating co-factor motifs (right). Each FOXA1 motif is either mutated (mut), orientation-reversed (rev), or converted into the strongest consensus (con). Both tables list the fold change and statistical significance of FOXA1 ChIP-ISO signal caused by these sequence variations. Two-tailed paired t-test. d) Example FOXA1 ChIP-ISO signals over CCND1e variants containing 0, 1, 2, or 3 original (ori) FOXA1 motifs or 1 consensus (con) FOXA1 motif. X: missing data. e) Box-and-whisker plots showing FOXA1 ChIP-ISO signals over otherwise identical CCND1e sequences containing WT or mutated AP-1, CEBPB, or SP1 motifs. Paired sequences are connected by a line. ****: p < 0.0001, ***: p < 0.001, **: p < 0.01, and ns: non-significant based on two-tailed paired t-test (same below, unless specified). f) Pearson correlation coefficient between FOXA1 ChIP-ISO signals and CCND1 sequence variables, calculated using the total set or the subset containing an AP-1 motif. These numbers reflect the level of impact of each variable on FOXA1 binding. g) FOXA1 ChIP-ISO signal as a function of the linear distance between the 3rd FOXA1 motif and the AP-1 motif. h) Relation between the FOXA1 ChIP-ISO signal and the total FOXA1 motif score over CCND1 variants ± AP-1 / CEBPB motifs. i) Effect of FOXA1 on AP-1 binding. Bar plots show FOXA1 and FOSL2 ChIP-qPCR for three biological replicates over integrated WT CCND1e or a variant with all three FOXA1 motifs mutated (Knock-in). ChIP-qPCR over a positive / negative control locus (P and N) and the native CCND1e are also shown. Error bars represent standard error (same below, unless specified).

To more accurately pinpoint sequence features affecting FOXA1 binding, we first replaced each FOXA1 motif with mutated, reversed, or consensus versions (**Extended Data Fig. 3c**). As expected, mutating/strengthening FOXA1 motifs significantly reduces/increases FOXA1 binding, respectively, while reversing their orientation has a minor effect (**Fig. 2c**). Consistent with **Fig. 2b**, the third FOXA1 motif is more influential than the other two (**Fig. 2c,d**). FOXA1 binding also increases with the number of FOXA1 motifs in a non-linear fashion (**Fig. 2d**), indicating that there is a synergistic effect among multiple adjacent binding sites.

We next examined the effect of other TFs that potentially co-bind with FOXA1. As there are eight co-factor motifs, the ChIP-ISO library includes all 256 combinations where each motif can be wild-type (WT) or mutated. We found that mutating AP-1 and CEBPB motifs leads to a significant decrease in FOXA1 binding (**Fig. 2c**). AP-1 has a particularly strong effect, as mutating its motif dramatically decreases FOXA1 binding to a level close to the background for almost all *CCND1e* variants (**Fig. 2e**). The presence of AP-1 highly correlates with FOXA1 binding, even more than the total FOXA1 motif score (**Fig. 2f**). Low-throughput FOSL2 (a subunit of AP-1) and FOXA1 ChIP confirmed the abolished binding of both factors when the AP-1 motif is mutated (**Extended Data Fig. 3d**). This data indicates that AP-1 is a crucial co-factor that potentiates FOXA1 binding to the *CCND1e*. Interestingly, both AP-1 and CEBPB motifs are immediately adjacent to the third FOXA1 motif, which has the largest impact on FOXA1 binding (**Fig. 2a,c**). To test the significance of this observation, we moved the AP-1 motif away from the third motif. We found that FOXA1 binding declines markedly with increasing distance (**Fig. 2g**), indicating that motif proximity is important for AP-1 facilitated FOXA1 binding.

Correlation analysis shows that FOXA1 binding is mostly affected by AP-1, its own motif, and CEBPB (**Fig. 2f**). To understand the interplay between these factors, we plotted the FOXA1 binding strength as a function of the total FOXA1 motif score in the presence or absence of AP-1 and CEBPB (**Fig. 2h**). Without AP-1 and CEBPB, FOXA1 can still bind strong motifs, but the presence of these two co-factors allows FOXA1 to target sub-optimal motifs, at least in the *CCND1e* context (**Fig. 2h**).

We also investigated the reciprocal relationship of FOXA1 on AP-1 binding to determine if AP-1 binds upstream of FOXA1 or if they bind cooperatively to enhance each other’s binding. We addressed this question by generating a new cell line containing the *CCND1e* with all three FOXA1 motifs mutated and measured AP-1 binding in the absence of FOXA1 with ChIP-qPCR. Notably, both FOXA1 and AP-1 binding are drastically reduced on this mutated *CCND1e* (**Fig. 2i**), supporting the scenario that the binding of these two TFs is mutually dependent and cooperative.

### AP-1 and CEBPB co-bind with FOXA1 and assist its binding genome-wide

The case study of the *CCND1e* demonstrates the importance of co-factors in FOXA1 binding. We next asked if this phenomena applies to other genomic loci and/or with other co-factors. We first evaluated the co-binding of FOXA1 with other TFs in A549 cells based on the overlap between their ChIP-seq peaks and the occurrence of their motifs in FOXA1 peaks (**Methods**). A large fraction (25% to 45%) of FOXA1 ChIP-seq peaks overlap with the peaks of AP-1 subunits JUNB, JUND, and FOSL2 (**Fig. 3a**), and vice versa (**Extended Data Fig. 4a**). Moreover, the most enriched motifs within FOXA1 peaks, aside from the FOXA1 motif itself, are those of the AP-1 subunits (*P*-value < 10^-1000^) (**Fig. 3a**). This data supports wide-spread co-binding of FOXA1 and AP-1. Many other TFs, including CEBPB, also display significant co-binding with FOXA1 (**Fig. 3a**).

**Fig 3:**
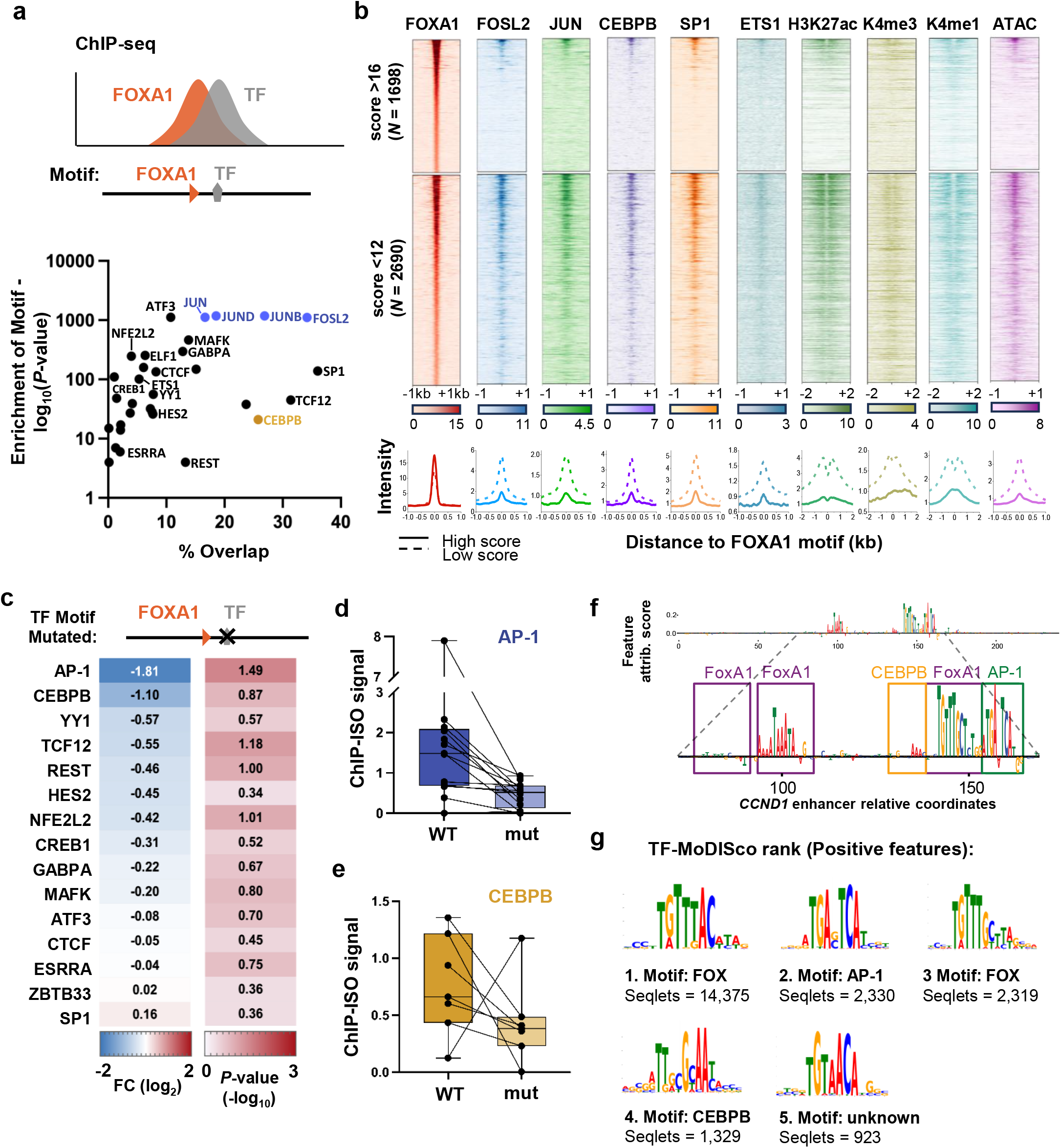
AP-1 and CEBPB co-bind with FOXA1 and assist its binding genome-wide. a) Bioinformatic analysis of FOXA1/TF co-binding. Top: Schematics of co-binding events. Bottom: for each TF, the dot plot shows the percentage of the overlapped FOXA1 ChIP-seq peaks (x axis) and the enrichment of its motif within FOXA1 peaks (y axis). TFs chosen for further analysis are labeled. AP-1 subunits are indicated in blue. b) Heatmaps of ChIP-seq / ATAC-seq and the corresponding intensity profiles in WT A549 cells over FOXA1 binding regions separated based on FOXA1 motif scores. Sequences in the top section contain strong consensus motifs (score >16), and the ones at the bottom contain weak motifs (score < 12). c) The effect on FOXA1 binding by mutating co-factor motifs. The ISO set used in panel c-e is derived from genomic sequences that show overlapped FOXA1 and TF ChIP-seq peaks in A549 and contain both motifs in proximity (<30bp). The table lists the fold change and statistical significance of FOXA1 ChIP-ISO signal upon mutations of co-factor motifs in these sequences. Two-tailed paired t-test. d & e) Box-and-whisker plots showing the changes of FOXA1 ChIP-ISO signal upon mutations of AP-1 (d) or CEBPB motif (e). f) DeepLIFT-Shap feature attribution scores highlighting features used by a sequence-trained CNN to predict FOXA1 binding in A549 cells at the CCND1e. g) Top 5 motifs detected by TF-MoDISco in the genome-wide DeepLIFT-Shap positive feature attribution scores at sites predicted by the CNN to be bound by FOXA1. The TF family of matching motifs is annotated for each motif, as is the number of seqlets used by TF-MoDISco to construct each motif.

Given that co-factors may permit FOXA1 binding at suboptimal motifs (**Fig. 2h**), we analyzed FOXA1-TF co-binding at FOXA1 sites with different motif strengths. We separated FOXA1 binding events near strong consensus motifs (scores > 16) or very weak ones (scores < 12) (**Fig. 3b**, left column). The average FOXA1 binding strength is comparable over these two sets of regions. Strikingly, co-binding predominantly occurs in the sites with low motif scores (**Fig. 3b**). These sites also show active histone marks and high chromatin accessibility (**Fig. 3b**). This data suggests that TF “hubs” tend to form over weaker motifs, and these co-binding events are more likely to be functional in gene regulation. This may represent a common strategy to ensure gene expression plasticity (see Discussion).

Co-binding between FOXA1 and TFs does not necessarily imply cooperativity among these factors. To test if the TFs identified in **Fig. 3a** indeed affect FOXA1 binding, we carried out additional mutational analyses for 15 TFs that show co-binding with FOXA1. For each TF, we selected 10-20 native genomic loci where it co-occupied by FOXA1 with proximal motifs (<30bp) (**Extended Data Fig. 4b**). The 193bp sequences from these loci, together with variants containing mutated TF motifs, were included in the ChIP-ISO library to evaluate the impact of these motifs on FOXA1 binding. AP-1 motif mutation again has the largest impact on FOXA1 binding, followed by CEBPB (**Fig. 3c-e**), indicating that these two factors promote FOXA1 binding at many genomic loci. These results also suggest that most co-localized TFs do not bind cooperatively with FOXA1.

To further assess whether co-factor motifs are predictive of genome-wide FOXA1 binding in A549 cells, we trained a convolutional neural network (CNN) to recognize FOXA1 ChIP-seq peaks using DNA sequence features (**Fig. 1a**). The CNN achieves high overall performance, with an area under precision-recall curve of 0.66 (calculated on held-out test sites). The CNN predictions of FOXA1 binding activities are generally consistent with ChIP-ISO measurements (**Extended Data Fig. 4c**). Using the DeepLift-SHAP feature attribution approach^38,39^, we characterized which DNA base positions contribute towards the CNN’s FOXA1 binding predictions at specific loci. Over *CCND1e*, for example, DeepLift-SHAP strongly highlights the second and third FOXA1 motifs and the AP-1 motif as positive contribution to FOXA1 binding (**Fig. 3f**). The feature attribution scores at the FOXA1, AP-1, and CEBPB motifs are weakened when mutated, and those from FOXA1 are strengthened when replaced with the consensus, consistent with ChIP-ISO measurements (**Extended Data Fig. 4d**). In addition, we ran the TF-MoDISco tool^40^ to compile feature attribution scores from across all ChIP-seq peaks into commonly occurring motif patterns. Alongside cognate FOXA1 binding motif variants, TF-MoDISco identifies the AP-1 and CEBPB motifs as the most prominent co-factor motifs that the CNN uses to predict FOXA1 binding (**Fig. 3g**). A GC-rich sequence similar to the SP1 motif is identified as the most negative feature (**Extended Data Fig. 4e**). In summary, our CNN analysis of FOXA1 ChIP-seq data is consistent with our findings that AP-1 and CEBPB assist FOXA1 binding genome-wide in A549 cells.

### AP-1 inhibition leads to motif-directed redistribution of FOXA1 binding in the genome

To further test the role of AP-1 in promoting FOXA1 binding, we measured the effect of knocking down AP-1 on genome-wide FOXA1 binding. Since the AP-1 family has multiple homologs that may have redundant functions, we took advantage of a dominant-negative protein A-FOS to inhibit global AP-1 binding. A-FOS dimerizes with JUN family proteins to form a heterodimer that cannot bind DNA^41,42^. We constructed an A549 cell line with doxycycline (Dox)-inducible A-FOS expression (**Fig. 4a**, **Extended Data Fig. 5a**). FOSL2 ChIP-seq verified that A-FOS expression leads to a near-complete inhibition of genome-wide AP-1 binding (**Fig. 4b**, **Extended Data Fig. 5b**). FOXA1 ChIP-seq in the ±Dox conditions shows that A-FOS induction causes significant reduction of FOXA1 binding over the sites where FOXA1 and AP-1 peaks overlap, while FOXA1 binding over non-overlapping sites remains unchanged (**Fig. 4b-d**, **Extended data Fig. 5c**). Differential binding analysis revealed 1,340 reduced and 234 enhanced FOXA1 peaks in the presence of A-FOS (“lost” vs “gained” peaks). Over 80% of the lost peaks contain AP-1 motifs and/or show AP-1 binding, much higher than the unchanged and the gained peaks (**Fig. 4e,f**). These results further support that AP-1 directs FOXA1 binding.

**Fig 4:**
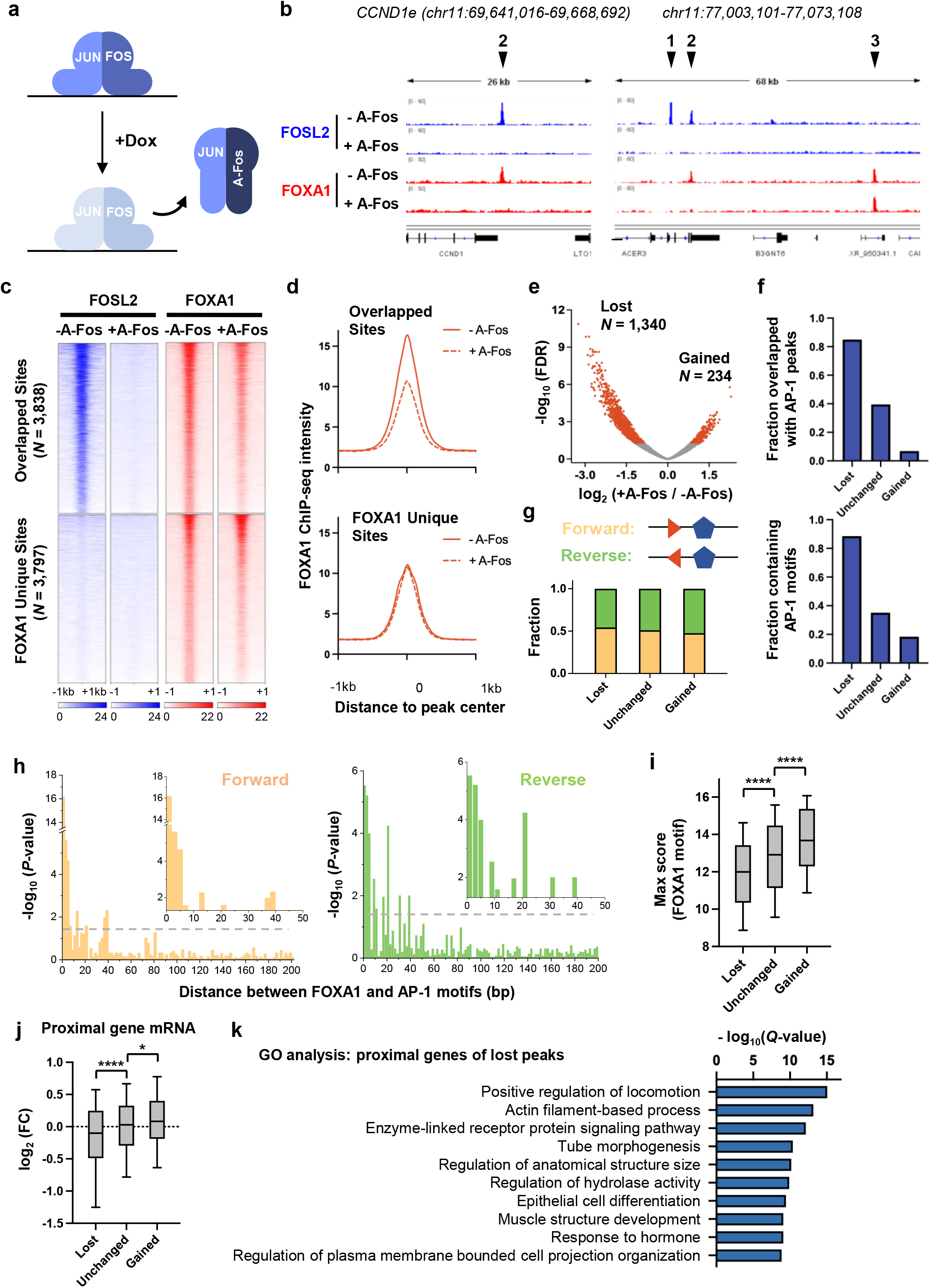
AP-1 inhibition leads to motif-directed redistribution of FOXA1 binding in the genome. a) Schematic showing the effect of A-FOS induction. Upon doxycycline-induced overexpression of A-FOS, it dimerizes with Jun and thus prevents Fos:Jun heterodimer formation and chromatin binding. b) Representative genomic tracks of FosL2 and FOXA1 ChIP-seq ± A-FOS induction in WT A549 cells. Arrows 1-3 demarcate examples of AP-1 unique, AP-1 / FOXA1 overlapped, and FOXA1 unique sites, respectively. In the presence of A-FOS, FOXA1 binding is significantly reduced at the overlapped site (2), but not the unique site (3). c) Heatmap of FosL2 and FOXA1 ChIP-seq signals over AP-1 / FOXA1 overlapped sites and FOXA1 unique sites ± A-FOS. d) Profiles of the average FOXA1 ChIP-seq intensities in panel c. e) Volcano plot showing differential FOXA1 binding ± A-FOS. Reduced (loss) and enhanced (gain) FOXA1 peaks in +A-FOS are highlighted. f) Overlap of loss / unchanged / gain FOXA1 peaks with AP-1. The upper panel shows the fraction that overlapped with FosL2 peaks, and the lower panel shows the fraction that contains AP-1 motifs. g) Distribution of the two orientations of FOXA1 / AP-1 motifs in loss / unchanged / gain peaks. Orange arrow: FOXA1 motif, blue pentagon: AP-1 (palindromic). h) Distribution enrichment of FOXA1 / AP-1 motif distances in loss peaks separated by the two orientations. Enrichment was calculated based on the histogram in Extended Data Fig. 5e using right-tailed two-proportion Z-test. i) The distributions of the maximum FOXA1 motif score per peak for loss / unchanged / gain FOXA1 peaks. j) Fold changes of the RNA-seq counts with A-FOS overexpression for the proximal genes near loss / unchanged / gain FOXA1 peaks. k) Top 10 enriched Gene Ontology (GO) terms of the proximal genes near the loss FOXA1 peaks.

We next analyzed whether AP-1-enhanced FOXA1 binding depends on specific configurations of their motifs, i.e. relative orientation and distance (**Extended Data Fig. 5d**). The two motif orientations are evenly distributed regardless of the peak category (**Fig. 4g**), while the two motifs are much more likely to be located within 8bp in the lost peaks (**Fig. 4h**, **Extended Data Fig. 5e**). These results, along with the data in **Fig. 2g**, show that proximity, but not a specific spacing or orientation between FOXA1 and AP-1 motifs, is required for their cooperativity. Weak enrichment is also observed near 10, 20, 30, and 40bp, suggesting that the rotational orientation of these two motifs on the same side of the DNA promotes cooperativity. In addition, we found that the maximum FOXA1 motif scores are significantly lower in lost peaks than in unchanged and gained peaks (**Fig. 4i**). This is consistent with the observation in **Fig. 3b** that FOXA1 binding over weaker motifs tends to be more AP-1 dependent. It also suggests that, upon AP-1 inhibition, FOXA1 is released from the weaker sites and re-distributed to stronger motifs.

To explore the functional role of FOXA1 binding events potentiated by AP-1, we conducted RNA-seq in cells ±A-FOS overexpression. We found that the genes proximal to lost peaks show significant down-regulation in the absence of AP-1, while the ones associated with gained peaks tend to be upregulated (**Fig. 4j**). Differential expression analysis also revealed the same trend (**Extended Data Fig. 6**). These results indicate that AP-1-facilitated FOXA1 binding events mostly mediate positive regulation of gene expression in A549 cells. Gene ontology (GO) analysis of the genes proximal to the lost peaks show enrichment in the cell migration, tissue development, and signal transduction categories (**Fig. 4k**), implying their cell-type-specific and differentiation-linked functions.

### *In vitro* study of FOXA1 binding and cooperativity with AP-1

To directly evaluate the intrinsic FOXA1 binding activity, we developed an electrophoretic mobility shift assay followed by sequencing (EMSA-seq) to measure the *in vitro* binding affinities between FOXA1 and all library sequences simultaneously (**Fig. 1a**; **Methods**). In this method, EMSA was performed using mixed library DNA incubated with purified FOXA1 at different concentrations (**Fig. 5a**, **Extended Data Fig. 7a**). Shifted (FOXA1-bound) vs unshifted (unbound) bands were then purified, PCR amplified, and subjected to amplicon sequencing. Normalized sequencing counts were converted into the “ratio bound *in vitro*” for each sequence, which was highly correlated among two replicates (**Extended Data Fig. 7b**; **Methods**).

**Fig 5:**
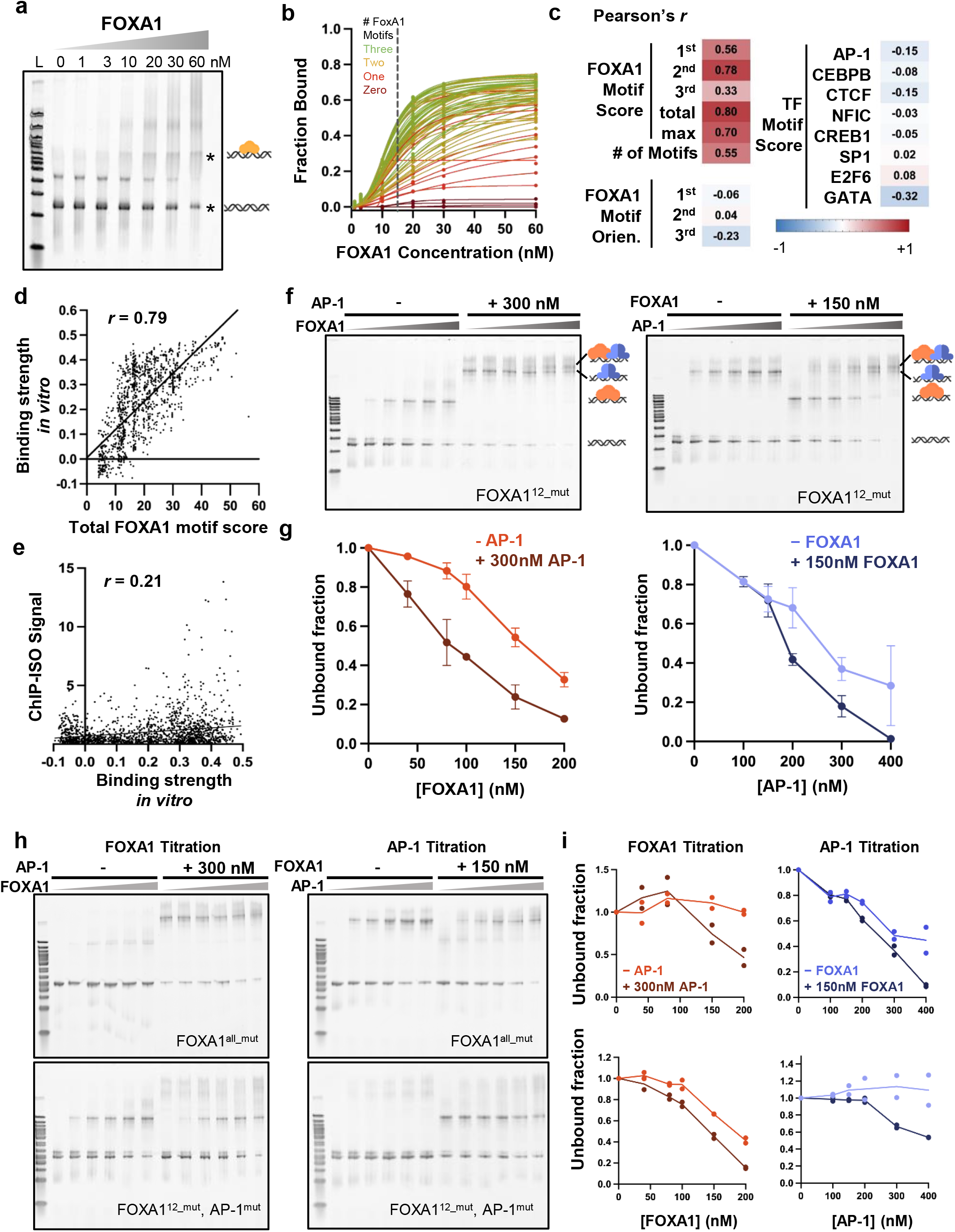
In vitro study of FOXA1 binding and cooperativity with AP-1. a) A representative EMSA-seq gel conducted on the ISO library with increasing levels of purified recombinant mouse FOXA1. The lower asterisk represents unbound oligonucleotides, and the upper asterisk indicates FOXA1-bound oligonucleotides. L: 100bp DNA ladder. b) FOXA1 bound fraction as a function of FOXA1 concentration measured by EMSA-seq. Data for a subset of the library, CCND1e variants, is plotted here. Green, yellow, orange, and red curves correspond to CCND1 variants with three, two, one, and zero FOXA1 motifs. The dotted line marks the fraction bound at 15 nM FOXA1, which is used to represent “b g strength in” in c-e. c) Pearson correlation coefficient between FOXA1 binding strength in vitro and CCND1 sequence variables. d) Correlation between the binding strength in vitro and total (summed) FOXA1 motif score for each sequence. e) Correlation between the binding strength in vivo (measured by ChIP-ISO) and that in vitro (EMSA-seq). f) Representative EMSA gels with FOXA1 titration ± AP-1 (left) or AP-1 titration ± FOXA1 (right) using a CCND1 variant with the first two FOXA1 motifs mutated (FOXA112_mut). Different populations are labeled on the right side of the gel (FOXA1 = orange, AP-1 = blue). g) Quantification of the EMSA gel in panel f. Error bar represents standard error for three replicates. h & i) Same as f & g but using different DNA templates and with two replicates performed on each template. Top: CCND1 variant with all three FOXA1 motifs mutated (FOXA1all_mut). Bottom: CCND1 variant with the first two FOXA1 motif and AP-1 motif mutated (FOXA112_mut, AP-1mut).

FOXA1 binding *in vitro* is primarily determined by the motif strength. Among the *CCND1e* variants, for example, FOXA1 binding generally increases with the number of motifs (**Fig. 5b**, **Extended Data Fig. 7c**). The EMSA-seq signals of the whole library are highly correlated with FOXA1 motif score (**Fig. 5c,d**). Different features of DNA shape play only a minor role (**Extended Data Fig. 7d**). Importantly, FOXA1 binding is no longer sensitive to mutations in AP-1 motifs *in vitro* (**Fig. 5c**, **Extended Data Fig. 7e**), confirming that the AP-1 effect on the FOXA1 ChIP-ISO signals is not due to inadvertent changes in intrinsic FOXA1 binding affinities. On the same library sequences, *in vitro* FOXA1 binding poorly correlates with that *in vivo* (**Fig. 5e**). This reinforces our previous finding that strong FOXA1 motifs are only partially responsible for FOXA1 binding *in vivo*.

To investigate potential cooperativity between AP-1 and FOXA1 *in vitro*, we purified recombinant AP-1 (**Extended Data Fig. 7a**) and performed low-throughput EMSAs with AP-1 and FOXA1 using *CCND1e* DNA. To focus on the co-binding between AP-1 and the most proximal FOXA1 motif, as indicated by **Fig. 2b,g**, we used a *CCND1e* template that has the first two FOXA1 motifs mutated (**Fig. 5f**). The gels show distinct bands for DNA bound by FOXA1 or AP-1 alone, and a super-shift for DNA bound by both factors (**Fig. 5f**). Quantification of the unbound band intensity shows that the presence of AP-1 moderately promotes the binding of FOXA1, and vice versa (**Fig. 5g**). Interestingly, we observed binding cooperativity between FOXA1 and AP-1 even in the absence of one factor’s motif (**Fig. 5h,i**). These results suggest that FOXA1 and AP-1 may exhibit protein-protein interactions that allow them to recruit each other without direct DNA binding. This can at least partially explain the interdependency and cooperativity of these two factors *in vivo*.

### FOXA1 binding is mostly determined by the local sequence, not the chromatin context

With the work above focusing on local sequences, we next explored how the larger-scale chromatin context can impact FOXA1 binding. In ChIP-ISO, native sequences containing FOXA1 motifs are moved from their endogenous loci to the euchromatic *AAVS1* site (**Extended Data Fig. 8a**). Comparison of FOXA1 occupancy at the native vs *AAVS1* locus therefore allows us to infer the effect from the endogenous chromatin.

We first applied this strategy to FOXA1 sites within euchromatic regions. We selected two sets of native sequences where FOXA1 binding cannot be explained by its motif strength: those with high-score FOXA1 motifs but mostly weak binding (set one) and vice versa (set two) (**Fig. 6a**, **Extended Data Fig. 8b**). EMSA-seq shows that most of these sequences are bound by FOXA1 *in vitro*, with slightly higher occupancies in set one (**Fig. 6b**). Strikingly, ChIP-ISO signals on these sequences at the *AAVS1* locus largely recapitulate the ChIP-seq intensities at the endogenous sites (**Fig. 6b**), indicating that FOXA1 binding are mostly determined by the local sequences. Such local signals again involve co-factors, with set two sites being more enriched with AP-1 and CEBPB motifs and showing higher AP-1 and CEBPB binding (**Fig. 6c**).

**Fig 6:**
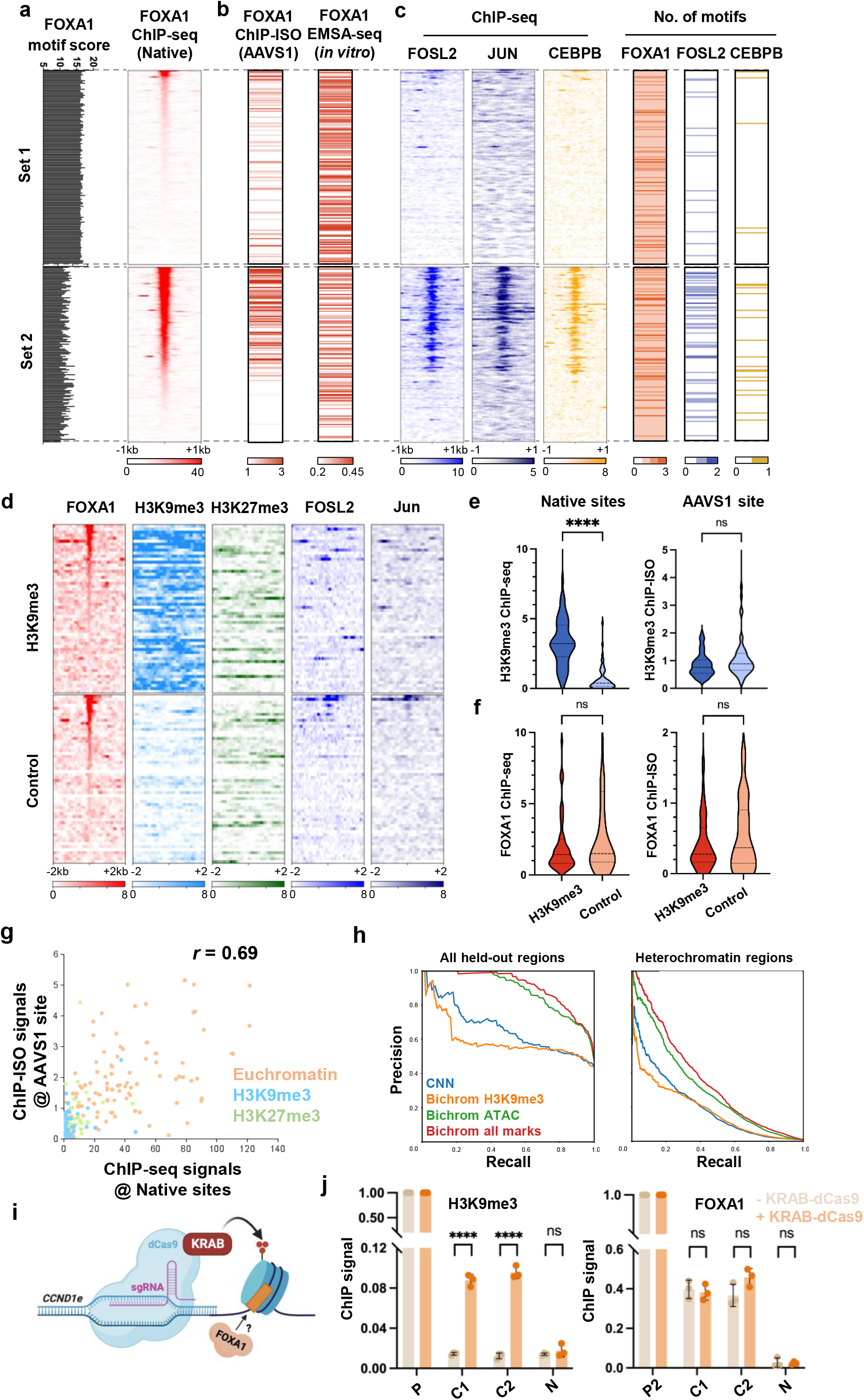
FOXA1 binding is mostly determined by the local sequence, not the chromatin context. a) ChIP-ISO test cases where strong / weak FOXA1 motifs show low / high FOXA1 binding (set 1 and 2, respectively). Left: FOXA1 motif scores. Right: heatmap of FOXA1 ChIP-seq signals in WT A549. b) Heatmap of FOXA1 ChIP-ISO signals (left) and EMSA-seq signals (fraction bound at 15 nM FOXA1, right) for sequences in panel a. c) Heatmap of FOSL2, JUN, and CEBPB ChIP-seq signals in WT A549 (left) and number of FOXA1, FOSL2, and CEBPB motifs for sequences in panel a. d) Heatmap of FOXA1, H3K9me3, H3K27me3, FOSL2, and Jun ChIP-seq signals in WT A549 for a set of ChIP-ISO library sequences derived from H3K9me3-marked regions (top) and a control set with comparable FOXA1 binding derived from euchromatic loci (bottom). e) Violin plot of H3K9me3 ChIP-seq signals for sequences in panel d at their native genomic loci (left) and ChIP-ISO signals of the same sequences at AAVS1 (right). f) Same as panel e, but for FOXA1. g) Correlation between FOXA1 ChIP-ISO and ChIP-seq signals for all sequences derived from the native genome in our library. Orange: sequences from euchromatin, blue: H3K9me3-marked heterochromatin, green: H3K27me3-marked heterochromatin. h) Precision recall curves showing performance of neural networks trained on FOXA1 ChIP-seq data in A549 cells. Each plot shows performance of a CNN trained using only sequence (blue lines); Bichrom trained using sequence and H3K9me3 (orange lines); Bichrom trained using sequence and ATAC-seq (green lines); and Bichrom trained using sequence and a selection of five histone marks (red lines). The left plot shows the performance of the neural networks across all held-out test sites, while the right plot shows performance at FOXA1 motif instances that overlap H3K9me3 or H3K27me3 peaks. i) Schematic of CRISPRi method where KRAB-dCas9 is induced by doxycycline to ectopically write H3K9me3 to the endogenous CCND1e in A549 cells. This system is used to measure the effect of H3K9me3 deposition on FOXA1 binding. j) H3K9me3 (left) and FOXA1 (right) ChIP-qPCR signals on three biological replicates at a positive control region (P), CCND1e (C1 and C2), and a negative control region (N) with (orange) or without (beige) KRAB-dCas9 induction. ****: p < 0.0001 and ns: non-significant based on two-tailed unpaired t-test.

We next investigated the effect of heterochromatin by selecting FOXA1 motifs from regions covered by H3K9me3 and H3K27me3. If heterochromatin has a strong inhibitory effect, we expect FOXA1 binding to increase when these sequences are transferred to euchromatin. We picked 56 sequences from H3K9me3-marked regions (**Fig. 6d**). For comparison, we assembled a control set in euchromatin that lacks H3K9me3 but exhibits matching levels of FOXA1 binding (**Fig. 6d-f**). The differences in H3K9me3 signals between these two sets of regions disappear after they are relocated to the *AAVS1* locus, confirming the elimination of H3K9me3 marks at the new site (**Fig. 6e**). However, this does not lead to enhanced FOXA1 binding, as the sequences originally from heterochromatin still exhibit the same FOXA1 binding levels as the euchromatic control (**Fig. 6f**). We performed the same experiments using FOXA1 sites from H3K27me3 regions and got similar results (**Extended Data Fig. 8c,d**). Combining the data from eu-and heterochromatin, FOXA1 binding at the *AAVS1* site is highly correlated with that in their native sites (*r* = 0.69) (**Fig. 6g**). Overall, this data suggests that the native chromatin context, including H3K9me3 and H3K27me3 marks, plays a minor role in FOXA1 binding.

To test whether H3K9me3 enables better prediction of FOXA1 binding genome-wide, we again turned to neural networks trained on FOXA1 ChIP-seq data. Specifically, we used our previously described Bichrom neural network architecture to integrate DNA sequence and various chromatin features into the training process^43^. Integrating ATAC-seq signals or a combination of histone marks into the neural network helps to improve performance in distinguishing held-out FOXA1-bound and unbound sites (**Fig. 6h**). However, integrating H3K9me3 alone alongside DNA-sequence features does not improve performance (**Fig. 6h**), suggesting that Bichrom is unable to learn any informative relationship between H3K9me3 and FOXA1 binding.

We noted that FOXA1 motifs in H3K9me3 covered regions are weaker and have no adjacent AP-1 motifs. To ensure that the absence of a heterochromatin effect is not simply due to the lack of suitable motifs, we artificially introduced H3K9me3 to a strong FOXA1 site by targeting KRAB-dCas9 to the endogenous *CCND1e* through CRISPRi (**Fig. 6i**). ChIP-qPCR shows robust H3K9me3 signal at the *CCND1e* upon 48 hours of KRAB-dCas9 induction compared to uninduced cells (**Fig. 6j**). Upon H3K9me3 deposition, we found that FOXA1 binding at the *CCND1e* does not significantly change compared to uninduced cells (**Fig. 6j**). These results are in line with our finding that native chromatin context plays only a minor role, if any, in FOXA1 binding. Together, we conclude that FOXA1’s binding specificity *in vivo* is more determined by the local sequence than the epigenetic background.

### Cell-type-specific binding of FOXA1 correlates with differential expression of AP-1

Previous studies have shown that FOXA1 has different binding patterns in different cell types^34^. Considering our findings above, we hypothesized that differential availability of co-factors may contribute to such cell-type specificity. We therefore analyzed the RNA-seq data of FOXA1 and AP-1 subunits in three cancer cell lines, A549, HepG2, and MCF-7. Interestingly, FOXA1 mRNA level is lower but AP-1 subunits are higher in A549 than the other two cell lines (**Fig. 7a**). Immunostaining confirmed this trend at the protein level (**Extended Data Fig. 9a**). Despite the lower expression of AP-1 in HepG2 and MCF-7, FOXA1 still co-binds with AP-1, but the level of overlap is significantly reduced compared to A549 (**Fig. 7b**).

**Fig 7:**
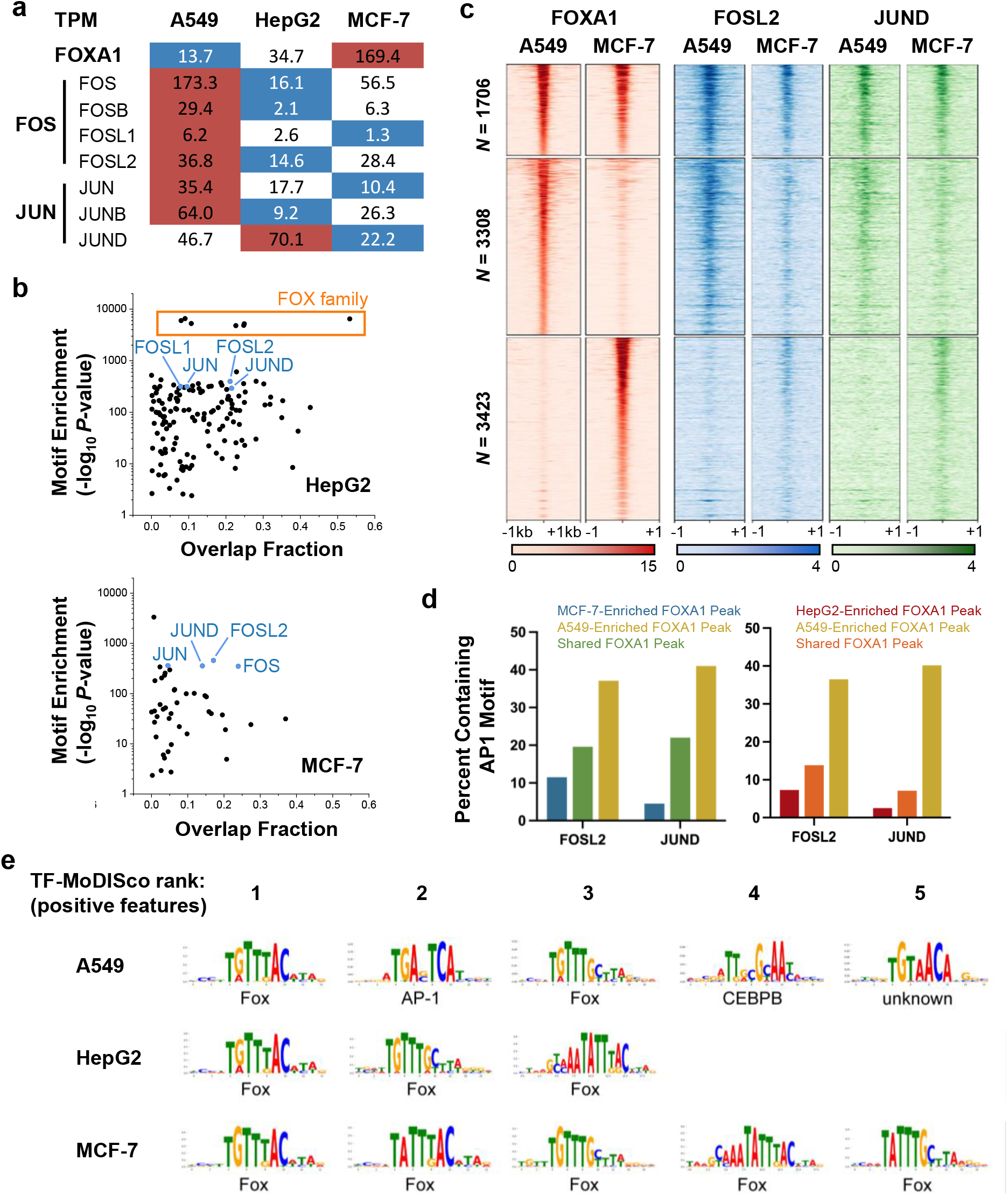
Cell-type-specific binding of FOXA1 correlates with differential expression of AP-1. a) RNA-seq counts, reported in transcripts per million (TPM), for FOXA1 and various AP-1 subunits in WT A549, HepG2, and MCF-7 cell lines. Red and blue mark the highest and lowest expression for each gene. b) Bioinformatic analysis of FOXA1 / TF co-binding in HepG2 (top) and MCF-7 (bottom). For each TF, the dot plot shows the percentage of the overlapped FOXA1 ChIP-seq peaks (x axis) and the enrichment of its motif within FOXA1 peaks (y axis). AP-1 subunits are indicated in blue. c) Differential FOXA1 binding analysis in A549 and MCF-7 cells, with heatmaps showing common (upper), A549-specific (middle), and MCF-7-specific (bottom) peaks. FOSL2 and JUND ChIP-seq signals over the same regions are shown on the right. d) Occurrence probability of FOSL2 or JUND motif in common or differential FOXA1 peaks. Left: A549 vs MCF-7; Right: A549 vs HepG2. e) Top ranking motifs detected by TF-MoDISco in the genome-wide DeepLIFT-Shap positive feature attribution scores at sites predicted by a sequence-trained CNN to be bound by FOXA1 in A549, HepG2, and MCF-7 cells. The top five ranking motifs are shown unless TF-MoDISco returned fewer than five motifs. The TF family of best-matching motifs is annotated for each motif.

The observations above raise the possibility that the abundance of AP-1 in A549 allows it to play a more dominant role in directing FOXA1 binding than in HepG2 and MCF-7. If this is true, we would expect a higher representation of AP-1 motifs in A549-specific FOXA1 peaks than the HepG2 or MCF-7-specific peaks. To test this prediction, we performed differential binding analysis of FOXA1 in A549 vs MCF-7/HepG2 and searched for motifs in common and cell-type specific peaks (**Fig. 7c**, **Extended Data Fig. 9b**). Indeed, A549-specific FOXA1 peaks are much more likely to contain AP-1 motifs, compared with shared peaks or MCF-7/HepG2-specific peaks (**Fig. 7d**). In A549 cells, AP-1 binding strongly correlates with that of FOXA1, and such correlation is weaker in the other two cell types (**Fig. 7c**). Mild AP-1 binding, however, is still detectable over MCF-7 and HepG2-specific FOXA1 sites, despite the lower enrichment of AP-1 motifs. This may be due to the tethering of AP-1 by FOXA1 as indicated by the *in vitro* binding assay (**Fig. 5h,i**). Overall, this data indicates that AP-1 is at least partially responsible for cell-type-specific binding of FOXA1.

To further characterize the importance of AP-1 in specifying cell-type-specific FOXA1 binding, we trained DNA sequence neural networks on 13 cell lines and tissue types, including the three above, with published FOXA1 ChIP-seq data (**Extended Data Fig. 10**). We again performed feature attribution analysis at ChIP-seq peaks and compiled informative patterns into motifs using TF-MoDISco. Consistent with the analysis above, AP-1 and CEBPB are not top features for promoting FOXA1 binding in MCF-7 and HepG2 (**Fig. 7e**). Interestingly, the AP-1 motif is identified as a highly informative feature for FOXA1 binding in the RT4 urinary bladder cell line, suggesting that AP-1 may assist FOXA1 binding in different cell types. In other cell lines, our neural networks identify additional informative co-factor motifs, including CTCF in 22Rv1 prostate carcinoma epithelial cells and AP-2 in both GP5d and SK-BR-3 cancer cell lines. We speculate that these factors may play analogous roles to AP-1 in assisting FOXA1 binding in these cell types.

## Discussion

Although PFs possess the unique ability to target nucleosomal motifs, they only bind a small subset of their motifs in higher eukaryotic genomes. Despite recent advances in genomic technology, extracting the genetic rules that govern TF/PF binding remains a formidable challenge. A significant hurdle arises from the fact that many genetic and epigenetic features can influence TF binding, and native genomes do not provide sufficient diversity to explore all possible combinations of these variables, especially within the constraints of evolution. In comparison, the ChIP-ISO assay utilizes artificially designed sequences^44^ that can circumvent evolutionary constraints and systematically perturb one genetic feature at a time. The synthetic sequences are inserted into the same genomic locus, which eliminates many variables caused by chromatin background, including the well-known ChIP artifact found near highly expressed genes^45^. Overall, the ChIP-ISO assay allows us to quantitatively dissect the contribution to TF binding from individual genetic features.

Cooperative binding is a commonly reported phenomenon among many TFs in mammalian cells, but PFs are often thought to function independently as the most upstream factors that interact with chromatin. FOXA1, for example, was shown to be able to associate with a high affinity motif embedded in a compact nucleosome array and open local chromatin *in vitro* without the assistance of other TFs or remodelers^31^. It is therefore surprising that co-factors are the main determinant of FOXA1 binding in A549 cells. These seemingly contradictory findings may be reconciled by considering the FOXA1 motif strength: while FOXA1 is able to bind to a subset of strong motifs in the absence of co-factors (**Fig 3b**), its binding on sub-optimal sites is strongly promoted by AP-1 and/or CEBPB (**Fig. 2h**, **Fig. 4i**). Co-binding with other TFs provides a mechanism for context-specific PF binding. Indeed, reduced AP-1 expression in MCF-7 and HepG2 cells releases FOXA1 near AP-1 motifs and allows it to occupy other genomic loci (**Fig 7**). This agrees with previous findings that the genomic distribution of FOXA1/2 can be affected by steroid receptors^46^, GATA4^7^, or PDX1^47^. These co-factor-dependent weak sites tend to be situated in open chromatin with active histone marks and are therefore more likely to carry regulatory functions (**Fig 3b**). Consistently, native enhancers often contain sub-optimal motifs with reduced TF binding affinities^48,49^. Overall, these findings suggest that, although FOXA1 may use consensus motifs to engage with chromatin by itself, weaker motifs in conjunction with co-factors may play a more functional role during development to generate cell-type-specific binding and regulation^47,50^.

AP-1 is a ubiquitously expressed TF that is highly represented in distal enhancers in many cell types^51^, where multiple TFs bind in hubs. It is therefore not surprising that the AP-1 motif is enriched near the binding sites of many TFs^51^. More relevant to this work, AP-1 was shown to co-bind with FOXA1 in breast and prostate cancer cells^34,52,53^ and with FOXA1/2 in pancreatic ductal adenocarcinoma^54^. In most of these cases, however, it is not clear if these factors bind independently, cooperatively, or hierarchically. Here, using well-controlled synthetic sequences, we clearly demonstrated that the binding of FOXA1 and AP-1 is mutually dependent. This finding, together with a previous proposal that AP-1 may also function as a PF^55^, raises an intriguing possibility that FOXA1 and AP-1 may bind together to achieve sufficient pioneering activities for the invasion and remodeling of closed chromatin in A549, which may contribute to cell-type-specific enhancer selection^56^.

The molecular mechanism of FOXA1 and AP-1 binding cooperativity requires further elucidation, but our current data provides some clues. First, the *in vitro* EMSA assay shows that AP-1 enhances FOXA1 binding, and vice versa. Such effects can even be observed on templates that lack the motif for one of the factors. These results indicate that there may be protein-protein interactions between these two factors that allow one to be tethered by the other. Consistent with this idea, weak AP-1 binding can be detected near many MCF-7 or HepG2 enriched FOXA1 binding sites, despite the fact that >90% of these sites lack the AP-1 motif (**Fig. 7c,d**). Second, AP-1 can stimulate FOXA1 binding with different distances and orientations between their motifs (**Fig. 4g,h**). This argues against a model where FOXA1 and AP-1 form a rigid complex with highly specific interactions. It is more likely that they have DNA-dependent weak and polymorphic interactions, and this type of “soft motif syntax” is commonly found among cooperative TFs^6,57,58^. Third, although the cooperativity between FOXA1 and AP-1 is detected *in vitro* on naked DNA, the effect is weaker than that *in vivo*. It is therefore possible that their cooperativity *in vivo* is also promoted by nucleosomes^59^. This would be consistent with the idea that FOXA1 and AP-1 may have co-pioneering activities.

Finally, our study suggests that the genomic background, including the heterochromatin marks, plays a minor role in FOXA1 binding. Heterochromatin has been reported in literature to both permit and inhibit PF binding^13,60,61^. For example, one study overexpressed a FOXA1 homolog, FOXA2, in immortalized foreskin fibroblasts cells and found that FOXA2 enrichment was generally depleted in H3K9me3 and H3K27me3 regions^7^. However, the same study also pointed out that, instead of a direct inhibitory effect from heterochromatin, this may be due to the lack of FOXA2 binding sites within these domains. Another study found that gain of FOXA2 binding at lamin-enriched sites is correlated with loss of H3K9me3, indicating its ability to target heterochromatin^62^. Here, we used two orthogonal methods, coupled with neural networks, to better understand the causal effect of heterochromatin on FOXA1 binding. Our ChIP-ISO results revealed that FOXA1 binds similarly to sequences in euchromatic vs heterochromatic context. Introducing ectopic H3K9me3 modifications over a strong FOXA1 site had no detectable effect on its binding. In addition, incorporation of H3K9me3 in the neural network analyses does not improve FOXA1 binding prediction. Together, these results suggest native chromatin context at most plays a minor role in FOXA1 binding. In summary, using the novel ChIP-ISO approach, in combination with *in vitro* and *in silico* analyses, our study demonstrated that cooperative binding with co-factors is the primary mechanism by which PFs achieve binding specificity. This result argues against the model that PFs exclusively function independently and explains the context-dependency of PF activities observed in multiple studies.

### Author Contributions

C.X., H.K., and L.B. conceived the experiments. C.X., H.K., and L.B. wrote and edited the manuscript with input from J.Y. and S.M. C.X. and H.K. designed the synthetic oligonucleotide library, performed ChIP-ISO and ChIP-seq experiments, and prepared and maintained cell lines. C.X. performed A-FOS-related experiments, RNA-seq, and immunofluorescence. H.K. performed *in vitro* and CRISPRi experiments. J.J. performed CRISPR genotyping and helped with A-FOS-related experiments. E.L. performed protein purifications. J.Y. built machine learning models and performed related analyses. C.X., H.K., and J.Y. performed bioinformatic analyses. L.B., S.M., and S.T. supervised and obtained funding for experiments.

## Methods

### Cell Lines

Wild-type A549 human lung carcinoma epithelial cells, a gift from Dr. Yanming Wang, were maintained in Ham’s F-12K (Kaighn’s) Medium (Gibco 21127022) supplemented with 10% FBS (Gibco 16000044) and 1% Penicillin-Streptomycin (Gibco 15070063). Wild-type HepG2 human liver cancer cells and MCF-7 human breast cancer cells were obtained from ATCC, and maintained in Dulbecco’s Modified Eagle Medium (DMEM) (Gibco 10569044) supplemented with 10% FBS (Gibco 16000044) and 1% Penicillin-Streptomycin (Gibco 15070063). All cells were cultured at 37°C in a humidified incubator with 5% CO_2_. Cells at passage number one were thawed and passaged at least an additional two times prior to experimental usage. All cell lines used in this study can be found in **Table S3**.

### Synthetic Oligonucleotide Design

The ChIP-ISO sequence library comprises 3,203 different sequences, each 229bp in length with 193bp variable regions and 18bp primer-binding regions on the two sides, each containing a BsaI or BbsI recognition site (different subsets of oligos use different primers and cutting sites) (**Extended Data Fig. 2a**). A detailed breakdown of the library composition can be found in **Table S1**. The synthetic oligo library was ordered from Agilent (Product #G7220A).

To design the ISO library with *CCND1e* variants in **Fig. 2**, a 193bp region from *CCND1e* (chr11:69,654,913-69,655,105) was selected (**Extended Data Fig. 3b**), and an internal BbsI cutting site was mutated to distinguish it from the native *CCND1e*. The library sequences were designed using a MATLAB program developed previously in the lab^63^. Each FOXA1 motif was mutated, reversed, or converted into consensus (**Extended Data Fig. 3c**). For the motifs of the other eight co-factors, the 1 to 3 most consensus bases were mutated to their complementary bases to maintain GC content, while avoiding interfering with neighboring motifs (checked by MEME “FIMO”^64^, Version 5.5.4). Some combinations of FOXA1 and co-factor motif variations were also included.

To design the ISO library containing native sequences with FOXA1 and TF cobinding in **Fig. 3**, overlaps between the top 30,000 genomic FOXA1 peaks from our FOXA1 ChIP-seq data and TF ChIP-seq peaks from ENCODE were identified using BEDTools “intersect intervals”^65^ (Version 2.30.0). The number of overlapping regions was divided by 30,000 (number of total FOXA1 peaks) to give the percent of FOXA1 peaks overlapping the TF. The enrichment of the TF motif within the 30,000 FOXA1 peaks was calculated using MEME “AME”^66^ (Version 5.5.4). A selection of TFs having a large percentage of FOXA1 peak overlaps and high motif enrichment in FOXA1 peaks were chosen for further analysis. FOXA1 and TF motifs within FOXA1 / TF overlapped regions were identified using FIMO with Position Weight Matrices (PWM) obtained from JASPAR or CIS-BP. These regions were filtered to select only for sequences containing a FOXA1 and TF motif within 8-30bp measured from the center of each motif. 10 to 20 regions were included in the synthetic library for each TF. Additionally, for each region, versions containing a mutated FOXA1 or TF motif were also included.

To design the ISO library “set 1” in **Fig. 6A**, which contains strong FOXA1 motifs but shows very weak or no FOXA1 binding, we first used FIMO to locate all FOXA1 motifs in the genome and calculated their motif scores based on PWM. A subset of motifs with score >16 and have no overlap with FOXA1 ChIP-seq peaks (evaluated by BEDTools “intersect intervals”) were selected for the library. For set2 where weak FOXA1 motifs are associated with strong ChIP-seq peaks, we first sorted FOXA1 ChIP-seq peaks based on their intensities using deepTools2^67^. Among the top 50% of the peaks, the corresponding genomic sequences (peak center +/-100bp) were retrieved using bedtools getfasta (Version 2.30.0), and FOXA1 motifs within these sequences were identified by FIMO. Sequences that contain a single motif with the score between 10-14.5 were selected for the library. To design the ISO library containing FOXA1 motifs covered by different epigenetic marks, FIMO was first used to identify FOXA1 motifs within FOXA1 ChIP-seq peaks and ChIP-seq peaks of different histone modifications. FOXA1 motifs present in both FOXA1 and H3K9me3/H3K27me3 peaks represent FOXA1-bound motifs within these repressive regions. FOXA1 motifs associated with FOXA1 peaks but absent in all histone modification datasets were labeled as FOXA1-bound motifs within unmarked regions. FOXA1-unbound motifs within repressive or unmarked regions were labeled similarly. A random subset of 50-60 sequences within each of these categories were included in the ChIP-ISO library, with the exception of FOXA1-bound motifs within H3K9me3-marked regions, in which all 21 regions were included. Each 193bp sequence was designed to include the genomic region surrounding the centered FOXA1 motif.

### Plasmid Construction

All plasmids and primers used in this study can be found in **Table S3**. The plasmid library backbone was derived from pAAVS1-Nst-MCS, which was a gift from Knut Woltjen (Addgene plasmid # 80487; http://n2t.net/addgene:80487; RRID:Addgene_80487). A ∼2kb human genomic region containing the *CCND1e* (chr11:69,653,809-69,655,876) was cloned between the PacI and SalI cutting sites. The two endogenous BsaI cutting sites on the resulting plasmid were mutated. The 193bp *CCND1e* sequence (chr11:69,654,913-69,655,105) was replaced by two BsaI cutting sites. The resulting plasmid named pCX1.10 was used as the backbone plasmid for ChIP-ISO plasmid library construction.

The plasmid expressing Cas9 and gRNA was derived from eSpCas9(1.1), a gift from Feng Zhang (Addgene plasmid # 71814; http://n2t.net/addgene:71814; RRID:Addgene_71814). sgRNA-T2 (5’-GGGGCCACTAGGGACAGGAT-3’) was cloned between the two BbsI cutting sites. The resulting plasmid was named pCX3.10.

To construct the plasmid for A-FOS overexpression, A-FOS sequence was PCR amplified from plasmid CMV500 A-FOS, a gift from Charles Vinson (Addgene plasmid # 33353; http://n2t.net/addgene:33353; RRID:Addgene_33353), and cloned into Xlone-GFP plasmid^68^ (a gift from Dr. Lance Lian), replacing the GFP gene between the KpnI and SpeI cutting sites. The resulting plasmid was named pCX4.1.

To design the piggyBac gRNA-containing vector used to randomly integrate *CCND1e*-targeting gRNAs into the genome, we followed a protocol described previously, with the following changes^69^. Two *CCND1e*-targeted gRNAs were designed using CHOPCHOP at a distance of roughly -300bp from the first FOXA1 motif and +300bp from the third FOXA1 motif. Oligonucleotides (IDT) corresponding to these two gRNA sequences were cloned into pGEP179_pX330K (Addgene 137882) and pX330S-2 (Addgene Kit 1000000055) according to the kit instructions. These gRNA-containing vectors were assembled by Gibson assembly into a single entry vector for Gateway cloning into pGEP163 (Addgene 137881), resulting in the plasmid pGEP163_CCND1_U2_D4.

### Generation of the Plasmid Library for ChIP-ISO

The synthetic oligonucleotide library was resuspended in TE buffer, pH 8.0, and diluted with water to 10 nM for PCR amplification. For each 1,000 types of oligonucleotides, 32 μl of 10 nM diluted synthetic library was amplified in a 400 μl PCR reaction (final template concentration 800 pM) for 13 cycles, using NEBNext Ultra II Q5 master mix (NEB M0544S). The PCR product was purified using Amicon Ultra-2mL 50K centrifugal filter (Millipore UFC205024) to exchange the PCR solution for 1X NEB CutSmart buffer. DNA concentration was estimated by agarose electrophoresis. ∼1.8 μg of the amplified library was digested with 60 U of BsaI/BbsI at 37°C overnight, followed by adding 30 U of extra BsaI/BbsI and digestion at 37°C for another two hours. The digestion products were purified with AMPure XP beads (Beckman Coulter MSPP-A63880) with a beads to DNA ratio of 1.8. 5 μg of pCX1.10 plasmid was digested with 120 U of BsaI at 37°C overnight, followed by adding 30 U of extra BsaI and digestion at 37°C for another two hours. 10 U of CIP was added into the digestion reaction, followed by incubation at 37°C for one hour to dephosphorylate 5’-ends. The linearized plasmid backbone was purified with E.Z.N.A. cycle pure kit (Omega D6492-02). 600 ng linearized plasmid backbone and 48 ng of digested library (molar ratio of 1:3) were ligated with 5 μL (2000 U) of T4 DNA ligase (NEB M0202S) in a 100 μL reaction. The ligation reaction was incubated at 16°C overnight, purified with E.Z.N.A. cycle pure kit and eluted with 30 μL of water. The purified ligation product was transformed into 5-alpha electrocompetent *E. coli* (NEB C2989) via electroporation. In each electroporation reaction, 25 μL of electrocompetent cells was transformed with 2.5 μL of purified ligation product. Adequate number of electroporation reactions were done to produce at least ∼100,000 colonies per 1,000 types of oligonucleotides. The *E. coli* cells were then pooled and grown overnight with ampicillin selection, followed by plasmid extraction using the E.Z.N.A. plasmid DNA maxi kit (Omega D6922-02).

### Generation of the Cell Library for ChIP-ISO

9.06x10^6^ wild-type A549 cells were plated per 15 cm dish in 22.5 mL of Ham’s F-12K (Kaighn’s) Medium supplemented with 10% FBS and 1% Penicillin-Streptomycin. 24 hours later, cells in each dish were transfected with 7.875 μg of pCX3.10 plasmid (expressing gRNA and Cas9), 23.625 μg of library plasmids, 96.75 μL of lipofectamine 3000 reagent (ThermoFisher L3000015) and 63 μL of P3000 reagent, which were diluted in Opti-MEM. 8-10 hours post-transfection, the media was replaced with fresh Ham’s F-12K (Kaighn’s) Medium supplemented with 10% FBS and 1% Penicillin-Streptomycin. 48 hours post-transfection, the cells in each dish were dissociated from the dish and splitted into two 15 cm dishes with Ham’s F-12K (Kaighn’s) Medium supplemented with 15% FBS, 1% Penicillin-Streptomycin and 600 μg/mL of G418. Media changes were performed every three to four days while the G418 selection was kept. When cell colonies were visible, the number of colonies was estimated by counting colonies inside randomly sampled grids on the dish under the microscope. Adequate number of transfection reactions were done, which produced ∼92,000 colonies for 3,203 types of sequences. The cells were then dissociated from the dishes, disaggregated, pooled and plated in new 15 cm dishes with Ham’s F-12K (Kaighn’s) Medium supplemented with 10% FBS, 1% Penicillin-Streptomycin and 300 μg/mL of G418. The pooled cell library was maintained and expanded for ChIP.

### Chromatin Immunoprecipitation (ChIP)

ChIP was performed with the cell library for ChP-ISO following a standard ChIP protocol. To fix protein-DNA interactions, formaldehyde (Ricca Chemical Company RSOF0010250A) was added to 2 X 10^7^ adherent log-phase cells to a final concentration of 1% and incubated for 10 minutes at room temperature. For FOXA1 and histone modification ChIP-ISO samples, 1.6 X 10^8^ cells and 8 X 10^7^ cells were fixed, respectively, in individual plates of 2 X 10^7^ cells for each replicate. Cross-linking was quenched by addition of glycine to a final concentration of 0.125 M and incubated for 5 minutes at room temperature. Cells were washed twice with cold 1X DPBS. Cells were scraped into cold 1X DPBS and pelleted at 4°C. In some cases, cell pellets were snap frozen using liquid nitrogen and stored at -80°C until ready to proceed. Fresh or thawed cell pellets were lysed by incubating cells in 2.5 mL cell lysis buffer (5 mM PIPES pH 8.0, 85 mM KCl, 0.5% NP-40) supplemented with protease inhibitor cocktail (Sigma-Aldrich P8340) for 10 minutes on ice. Cell nuclei were pelleted at 4°C and lysed in 150 μL nuclei lysis buffer (50 mM Tris-Cl pH 8.0, 10 mM EDTA, 1% SDS) supplemented with protease inhibitor cocktail for 10 minutes on ice. Chromatin was fragmented in Diagenode Pico with a circulating water bath at 4°C using the Shear and Go Easy Mode setting for 3 cycles (30 seconds on followed by 30 seconds off). Sonicated chromatin was centrifuged to remove cell debris and residual SDS precipitate. The supernatant containing sheared chromatin was pooled across all replicates, a 50 μL input DNA sample was reserved, and the remaining pool was split again into eight (FOXA1) or four (histone modification) chromatin samples. In some cases, supernatant containing sheared chromatin was snap frozen using liquid nitrogen and stored at -80°C until ready to proceed.

For each ChIP sample, 20 μL Magna ChIP™ Protein A+G Magnetic Beads (Sigma-Aldrich 16-663) was washed four times with 1X DPBS supplemented with 5 mg/mL BSA and subsequently crosslinked to 5 μg antibody (Anti-FOXA1 antibody: GeneTex, GTX100308; Anti-Fra2 antibody: Cell Signaling Technology, 19967S; Anti-H3K9me3 antibody: abcam, ab8898; and Anti-H3K27me3 antibody: abcam, ab6002) for two hours at 4°C. Antibody-crosslinked magnetic beads were washed four additional times with the DPBS/BSA solution. Each chromatin sample except the input was incubated with the washed antibody-crosslinked magnetic beads for two hours at 4°C. Next, the magnetic beads were washed five times with LiCl wash buffer (100 mM Tris pH 7.5, 500 mM LiCl, 1% NP-40, 1% sodium deoxycholate) and once with 1X TE buffer (10 mM Tris-HCl pH 7.5, 0.1 mM Na_2_EDTA) at room temperature. Immunoprecipitated chromatin was eluted from the magnetic beads by incubating in IP elution buffer (1% SDS, 0.1 M NaHCO_3_) for 1 hour at 65°C. The collected supernatant (containing immunoprecipitated chromatin) and the reserved input sample were both incubated with 40 μg RNase A and NaCl to a final concentration of 0.37 M overnight at 65°C. The next day, 80 μg proteinase K was added to the ChIP sample and 400 μg was added to the input sample and both were incubated at 55°C for 2 hours. The ChIP and input DNA was purified via phenol-chloroform extraction. At the DNA elution step, the individual ChIP DNA pellets were resuspended together to achieve a more concentrated sample.

Prior to any downstream applications, the success of the ChIP reaction was determined via qPCR using various positive, negative, and locus-specific primer pairs. qPCRs were performed using the Agilent AriaMx Real-Time PCR System with the SYBR Green optical module (Emission 516.0 nm, Excitation 462.0 nm). For ChIP-seq, the same ChIP protocol was used, but with the following modifications. For each ChIP biological replicate, only 2 X 10^7^ adherent log-phase cells were fixed. 50 μL of sheared chromatin was reserved per replicate as the genomic input sample. At the DNA elution step, each ChIP pellet was resuspended separately.

### Amplicon Sequencing for ChIP-ISO

To amplify the integrated ChIP-ISO library sequences while excluding other genomic DNA, including native *CCND1e*, we performed two rounds of PCR amplification with a BbsI digestion step in between (**Extended data Fig. 2h**). The primer pairs for the first round of PCR contain regions annealing to the *CCND1e* (outside the 193bp library sequence) at the 3’-ends, partial Illumina TruSeq adaptor sequences at the 5’-end and 0-3 random nucleotide spacers in between to increase sequence complexity. Primers with different numbers of spacers are mixed in equimolar ratio for the first round of PCR. Preliminary PCR tests were performed to decide the optimal cycle number that keeps the PCR reactions in exponential phase. We used 23-25 cycles for our first round of PCR. For the first round of PCR, 30 μl of ChIP DNA was amplified in a 100 μl PCR reaction using NEBNext Ultra II Q5 master mix. The PCR products were purified with AMPure XP beads with a beads to DNA ratio of 0.9. 15 μL of the purified PCR product was digested with 20 U of BbsI in a 30 μL reaction at 37°C for two hours. The digestion products were purified with AMPure XP beads with a beads to DNA ratio of 0.9, followed by the second round of PCR. The primer pairs for the second round of PCR contain the rest of the Illumina TruSeq adaptor sequences and sample indexes. For the second round of PCR, 2 μl of purified digestion product was amplified in a 50 μl PCR reaction for 8 cycles using NEBNext Ultra II Q5 master mix. The PCR products were purified with AMPure XP beads with a beads to DNA ratio of 0.8. Quality control was conducted with TapeStation (Agilent). 30 million paired-end 150bp reads were obtained for each ChIP-ISO and input sample using a NextSeq 2000 (Illumina) instrument. Demultiplexing was performed using DRAGEN BCL Convert (v3.8.4).

### Sequencing Data Analysis for ChIP-ISO

Raw sequencing reads were filtered using fastp with default settings^70^ (Version 0.23.2), and the first three nucleotides were trimmed from each read using cutadapt^71^ (Version 4.4) to remove the 0-3 random nucleotide spacers introduced by the amplicon primers. Processed forward and reverse reads were merged into single reads based on their overlapping regions using NGmerge^72^ (Version 0.1). Merged reads were aligned to a FASTA file containing all ChIP-ISO library sequences (including their reversely ligated versions) using BWA-MEM2^73^ (Version 2.2.1). BAM alignments containing at least 2 mismatched nucleotides were filtered out using BAMtools^74^ (Version 2.4.0). The number of filtered reads aligning to each ChIP-ISO library sequence was counted for each ChIP and input sample and normalized to the total number of sequencing reads for each sample. ChIP-ISO signal was calculated by dividing the normalized number of ChIP counts by the normalized number of input counts. Any sequence having fewer than 1000 input counts was excluded from further analyses, resulting in FOXA1 ChIP-ISO signals for 1,882 sequences. ChIP-ISO signal is reported and plotted as the average of two independent biological ChIP-ISO replicates.

### Low-throughput ChIP-qPCR test of binding on single integrated sequences

The overall process was the same as ChIP-ISO. Instead of the synthetic oligonucleotide library, single synthetic sequences were cloned into the pCX1.10 plasmid backbone. Wild-type A549 cells were transfected with the resulting plasmids individually, together with pCX3.10 plasmid. For each synthetic sequence, the cell colonies were pooled together after G418 selection, and expanded for ChIP. To measure TF binding to the integrated synthetic sequences and the native *CCND1* enhancer separately, quantitative PCR (qPCR) was conducted with primer pairs that can distinguish between the integrated and the native sequences (**Table S3**). Locked nucleic acids (LNAs) were incorporated into the primers to increase specificity.

### ChIP-seq and Data Analysis

Sequencing library was constructed by NEBNext ultra II DNA library prep kit (NEB E7103L). 50 million paired-end 50bp reads were obtained for each ChIP and input sample using a NextSeq 2000 instrument. Paired-end reads were filtered using fastp with default settings and subsequently aligned to the human reference genome (hg38) using BWA-MEM2. Resulting BAM files were filtered for MAPQ scores > 20 using SAMtools^75^ (Version 1.8). Mapped regions within the ENCODE Blacklist were excluded from further analysis^76^ (hg38 Version 2). Read coverage was obtained separately for input and ChIP samples using deepTools “bamCoverage”, with a bin size of 10bp^67^ (Version 3.5.1), and visualized in IGV (Version 2.8.12) or the UCSC Genome Browser. ChIP peaks were called from pooled ChIP replicates using MACS2 callpeak with the default settings^77^ (Version 2.1.1.20160309). Heatmaps and intensity profiles were generated using computeMatrix, plotHeatmap, and plotProfile functions in deepTools2^67^ (Version 3.5.4).

### RNA-seq

Cells were lysed by Trizol (ThermoFisher 15596026), extracted by 0.2 volume of chloroform, followed by adding equal volume of 100% ethanol. RNeasy kit (Qiagen 74104) was then used to purify the RNA. 3 µg of purified RNA was treated with 2 U of RNase-free DNase I, and purified again with RNeasy kit. Sequencing library was constructed by the Illumina Stranded mRNA Prep kit (Illumina 20040532). 30 million paired-end 50bp reads were obtained for each RNA-seq sample using a NextSeq 2000 instrument. Data analysis was conducted based on a protocol from Batut et al. 2021^78^.

### A-FOS-Related Experiments and Data Analysis

To construct A549 ePB tet-on A-FOS cell line, 6x10^5^ wild-type A549 cells were plated in a 6-well plate well in Ham’s F-12K (Kaighn’s) Medium supplemented with 10% FBS and 1% Penicillin-Streptomycin. 24 hours later, cells were transfected with 0.72 μg of piggyBac transposase plasmid (System Biosciences PB210PA-1) and 1.78 μg of pCX4.1 plasmid using Lipofectamine 3000 transfection reagent. 12 hours post-transfection, the media was replaced with fresh medium. 48 hours post-transfection, the cells in the well were dissociated from the dish and splitted into two 6-well plate wells with Ham’s F-12K (Kaighn’s) Medium supplemented with 15% FBS, 1% Penicillin-Streptomycin and 10 μg/mL of blasticidin. Media changes were performed every three to four days while the blasticidin selection was kept. When cell colonies were visible, the cells were then dissociated from the wells, disaggregated, pooled and maintained in Ham’s F-12K (Kaighn’s) Medium supplemented with 10% FBS, 1% Penicillin-Streptomycin and 5 μg/mL of blasticidin. The expanded cells were subject to immunofluorescence, ChIP-seq and RNA-seq.

To identify lost, unchanged and gained FOXA1 ChIP-seq peaks in +Dox versus-Dox conditions, differential binding analysis was conducted with Bioconductor “DiffBind”^79^ (Version 3.18) using default settings. The regions within each category were converted from BED to FASTA format using BEDTools “GetFastaBed”, and FOXA1 and AP-1 motif scanning was performed inside these regions using MEME “FIMO”. To identify proximal genes of FOXA1 ChIP-seq peaks, MEME “T-Gene”^80^ (Version 5.5.4) was first used to predict target genes for each category of peaks. The predicted target genes, whose distances to the corresponding ChIP-seq peaks are smaller than 100 kb, are selected as the proximal genes for further analysis. Gene ontology (GO) analysis on the proximal genes of the lost peaks was performed by “Metascape”^81^ (Version 3.5.20230501) using default settings.

### Recombinant Protein Expression and Purification

Mouse FOXA1 (UniProtKB: P35582) fused to an N-terminal 6x-histidine tag was expressed in BL21(DE3)pLysS *E. coli* cells (Novagen 69388-3) at 37°C for 3 hours using the bacterial expression plasmid pET-28b-FOXA1, a gift from K. Zaret, University of Pennsylvania Perelman School of Medicine, Philadelphia, PA^82^. Harvested bacterial cells were resuspended and lysed by sonication in P300 buffer (50 mM sodium phosphate pH 7.0, 300 mM NaCl, 5 mM 2-mercaptoethanol, 1 mM benzamidine). Following extraction of soluble proteins, insoluble material containing FOXA1 was resuspended and sonicated in P300 buffer with 7 M urea added. Solubilized FOXA1 was isolated using Ni-NTA chromatography (GoldBio H-350-25), and further purified by Source S cation-exchange chromatography (Cytiva 17-0944-01). FOXA1 protein was stored in 8 mM HEPES pH 7.5, 80 mM NaCl, 8 mM 2-mercaptoethanol, 0.7 M urea, 20% glycerol.

Genes encoding human c-Fos (UniprotKB: P01100) fused to an N-terminal 6x-histidine tag and untagged human c-Jun (UniprotKB: P05412) were subcloned into pST39^83^ and pST50Tr^84^, respectively, from pST39-F:cJun/6xHis:cFos, a gift from C.M. Chiang, UT Southwestern Medical Center, Dallas, TX^85^. Expression of each protein was carried out separately in Rosetta2(DE3)pLysS *E. coli* cells (Novagen 70951) at 37°C. Soluble proteins were extracted as described for FOXA1, and insoluble materials containing c-Fos and c-Jun were processed separately. c-Fos was solubilized by resuspension and sonication in T100 buffer (20 mM Tris-HCl pH 7.5, 100 mM NaCl, 5 mM DTT). Following centrifugation, clarified extract was dialyzed into P300 buffer containing 7 M urea, and c-Fos was partially purified from it using Ni-NTA chromatography. Insoluble material containing c-Jun was washed three times in T100 buffer, followed by solubilization in 20 mM Tris-HCl pH 7.5, 1 mM EDTA, 1 mM DTT, 6 M guanidine-HCl. Refolding of c-Fos/c-Jun heterodimers was performed by stepwise dialysis as described by Ferguson and Goodrich, 2001^86^ and purified by cobalt metal-affinity chromatography (Talon resin, Clontech 635652). cFos/cJun was stored in 18 mM Tris-HCl pH 7.5, 90 mM NaCl, 9 mM 2-mercaptoethanol, 20% glycerol. Proteins were analyzed by SDS-PAGE (**Extended Data Fig. 6a**).

### Electrophoretic Mobility Shift Assay with Sequencing (EMSA-seq)

A pooled equimolar mixture of the ChIP-ISO synthetic oligonucleotide library was PCR-amplified and purified via agarose gel purification (Thermo Scientific K0691). The EMSA protocol was adapted from Garcia et al. 2019^78^. Briefly, 100 nM ChIP-ISO synthetic oligonucleotide library was incubated with 50X non-specific competitor DNA and 0-60 nM recombinant mouse 6xHis-FOXA1 in binding buffer (10 mM Tris-HCl pH 7.5, 1 mM MgCl_2_, 10 µM ZnCl_2_, 50 mM KCl, 3 mg/mL BSA, 10% glycerol, and 1 mM DTT) at room temperature for 30 minutes. Free and FOXA1-bound library sequences were separated on a 7.5% non-denaturing polyacrylamide gel (Bio-Rad 4561026) run in 1X Tris-Glycine at 200V at room temperature for 30 minutes. Gels were stained with 1 µg/mL Ethidium Bromide (Invitrogen 15585011) in 1X Tris-Glycine for 10 minutes at room temperature. Stained gels were visualized with a Bio-Rad GelDoc Go Imaging System using the Ethidium Bromide setting (**Fig. 5a**). The FOXA1-bound and-unbound DNA bands were excised at each FOXA1 concentration, and the DNA was eluted from each polyacrylamide gel slice following a User-Developed Protocol for extraction of DNA fragments from polyacrylamide gel using the QIAGEN QIAquick Gel Extraction Kit (QIAGEN 28704). The gel-extracted DNA was PCR-amplified and purified using AMPure XP Beads (Beckman Coulter A63880) with a 0.9x bead cleanup ratio. The library was created in the same manner as the ChIP-ISO library.

30 million paired-end 150bp reads were obtained for FOXA1-bound and-unbound DNA samples at each protein concentration using a NextSeq 2000 instrument. The sequencing data was processed and analyzed in the same manner as the ChIP-ISO datasets. The number of filtered paired-end reads aligning to each ChIP-ISO library sequence was counted for each bound and unbound sample and normalized to the number of sequencing reads for each sample. The ratio of FOXA1-bound DNA to total input (FOXA1-bound + FOXA1-unbound) DNA was calculated for each synthetic oligonucleotide library sequence at each FOXA1 concentration and normalized to the corresponding FOXA1-bound ratio of negative control CCND1e-FOXA1^all_mut^ (Index: 22). The resulting negative-normalized FOXA1-bound ratios were further normalized to the highest ratio across all FOXA1 concentrations, forcing ratios to fall between 0-1 for ease of analysis. For simplicity, these values are called the “ratio bound *in vitro*.” To correct for systematic error, values from the first EMSA-seq replicate were adjusted such that the two replicates approximated *r* = 1 (**Extended Data Fig. 7b**). To determine the FOXA1 concentration at which the ratio bound *in vitro* falls into the linear range for most sequences, these values were plotted for all library sequences in the top-down *CCND1e* FOXA1 motif category (**Fig. 5b**). For each sequence, the data points were fit with the Hill slope equation Y=B_max_*X_h_/(K_dh_ + X_h_), where X = FOXA1 concentration and Y = ratio bound *in vitro*, to model specific FOXA1 binding. For all quantitative analyses, the ratio of each synthetic oligonucleotide library sequence bound by FOXA1 *in vitro* was approximated by its FOXA1-bound ratio at 15 nM FOXA1, which was extrapolated by averaging its FOXA1-bound ratios at 10 nM and 20 nM FOXA1. This FOXA1 concentration was chosen because it falls into the linear range of the FOXA1 binding curves of most sequences (**Fig. 5b**).

### FOXA1 and AP-1 Co-Binding Electrophoretic Mobility Shift Assays (EMSAs)

Three *CCND1e* DNA templates were designed to include different FOXA1 and AP-1 motif mutants. FOXA1^12_mut^ has mutations in the two upstream FOXA1 motifs, FOXA1^all_mut^ has mutations in all three FOXA1 motifs, and FOXA1^12_mut^, AP-1^mut^ has mutations in the two upstream FOXA1 motifs and single AP-1 motif. These DNA templates were individually PCR-amplified and purified using a PCR clean-up kit (Omega Bio-tek D6492). In the FOXA1 EMSAs, 100 nM *CCND1e* DNA was incubated with 50X non-specific competitor DNA and 0-200 nM recombinant mouse 6xHis-FOXA1 in DNA-binding buffer (10 mM Tris-HCl pH 7.5, 1 mM MgCl_2_, 10 µM ZnCl_2_, 50 mM KCl, 3 mg/mL BSA, 10% glycerol, and 1 mM DTT), with or without 300 nM recombinant human AP-1. Similarly, in the AP-1 EMSAs, up to 400 nM cJun/6His:cFos was titrated into the same buffer/DNA solution, with or without 150 nM FOXA1. The EMSA samples were incubated at room temperature for 30 minutes and separated on a 7.5% non-denaturing polyacrylamide gel (Bio-Rad 4561026) run in 1X Tris-Glycine at 200V at room temperature for 30 minutes. Gels were stained with 1 µg/mL Ethidium Bromide (Invitrogen 15585011) in 1X Tris-Glycine for 10 minutes at room temperature. Stained gels were visualized with a Bio-Rad GelDoc Go Imaging System using the Ethidium Bromide setting (**Fig. 5f** & **h**). For the FOXA1 titration EMSAs, the unbound fraction was calculated at each FOXA1 concentration by normalizing the intensity of the free band to the intensity of the free band at 0 nM FOXA1 ± AP-1 (**Fig. 5g** & **i**). The equivalent calculations were performed for the AP-1 titration EMSAs.

### CRISPRi

To integrate the KRAB-dCas9 construct into the AAVS1 locus, 2 X 10^6^ cells were co-transfected with 10 µg pT077 (Addgene 137879), 1.5 µg AAVS1 TALEN L (Addgene 59025) and 1.5 µg AAVS1 TALEN R (Addgene 59026) using Lipofectamine 3000 Transfection Reagent (Invitrogen L3000015) according to the manufacturer’s protocol. Transfected cells were transferred to a medium supplemented with 700 μg/mL G418 (Gibco 10131035) 24 hours post-transfection and maintained in this medium to allow for single-cell colony formation (∼14 days). Single colonies were picked and seeded into 24-well plates. Colonies were maintained in G418-supplemented medium until they reached sufficient cell density, and those that retained normal cell morphology and growth rate were split for visualization of EGFP and maintenance. To visualize the EGFP expression of each colony, cells were plated into two wells of an 8-well dish (ibidi 80806); one well of cells was induced with 1 μg/mL doxycycline (Sigma-Aldrich D5207) 24 hours after plating and one well was left untreated. 48 hours post-induction, inducible expression of KRAB-dCas9 was confirmed by measuring EGFP expression in induced cells normalized to untreated cells. Colonies with high relative EGFP expression and homogeneity were frozen for storage. The colony with the highest EGFP expression and homogeneity was validated using genotyping PCR to confirm KRAB-dCas9 integration at the AAVS1 locus. To randomly integrate *CCND1e*-targeting gRNAs throughout the genome of KRAB-dCas9 cells, 6 X 10^5^ cells were co-transfected with 5 µg of gRNA-containing piggyBac vector and 1 µg of piggyBac transposase plasmid (System Biosciences PB210PA-1) using Lipofectamine 3000 Transfection Reagent. Transfected cells were transferred to a medium supplemented with 700 µg/mL G418 and 10 µg/mL Blasticidin S HCl (Gibco A11139) 24 hours post-transfection and maintained in this medium to allow for single-cell colony formation (∼14 days). Constitutive expression of gRNAs was confirmed by measuring the expression of mRFP in a mixed cell population. Anti-Histone H3 (tri-methyl K9) antibody (abcam ab8898) was used to perform H3K9me3 ChIP on the mixed cell population, followed by qPCR to confirm H3K9me3 deposition at the *CCND1e*. Anti-FOXA1 antibody (GeneTex GTX100308) was used to perform FOXA1 ChIP on the mixed cell population, followed by qPCR, to monitor any changes to FOXA1 binding at the *CCND1e* upon H3K9me3 deposition.

### Cell Type-Specific FOXA1 and AP-1 Motifs, Binding, and Expression

To identify FOXA1 binding sites that are enriched, shared, or depleted in A549 cells compared to HepG2 / MCF-7 cells, a HepG2 FOXA1 ChIP-seq BED file containing significant peaks from at least two replicates was acquired from the ENCODE Database, sorted by coordinate using BEDTools “sortBED”, and pooled with the top 30,000 significant WT A549 FOXA1 ChIP-seq peaks from two replicates we acquired. To determine the number of reads aligning to each region in the pooled BED file across cell types and replicates, the pooled BED file and the corresponding BAM files from two HepG2 and two A549 replicates were input into BEDTools “MultiCovBed” using default parameters. The output from this tool was input into Bioconductor “edgeR”^87^ (Version 3.34.0) using default settings to identify regions enriched, shared, or depleted in FOXA1 binding in A549 vs HepG2. Regions with a log_2_ fold-change less than -1 were labeled A549-depleted, between -1 and 1 were labeled shared, and greater than 1 were labeled A549-enriched. The regions within each category were converted from BED to FASTA format using BEDTools “GetFastaBed”, and each category was input into MEME “FIMO” to identify the number of regions containing FOS or JUN motifs. The percent of regions in each category containing each individual FOS or JUN motif was calculated.

### Immunofluorescence

Immunofluorescence experiments were performed according to a protocol from Yoney et al. 2022^88^. The following primary antibodies and dilutions were used: FLAG (mouse monoclonal, Millipore-Sigma, F1804, 1:1000), FOXA1 (rabbit polyclonal, GeneTex, GTX100308, 1:500), and FOSL1 (mouse monoclonal, Santa Cruz Biotechnology, sc-28310, 1:50). The following secondary antibodies and dilutions were used: goat anti-mouse IgG(H+L) (Alexa Fluor 594, ThermoFisher, A-11005, 1:1000), and goat anti-rabbit IgG(H+L) (Alexa Fluor 488, ThermoFisher, A-11008, 1:500).

Immunostained A549 ePB tet-on A-FOS cells were imaged using Leica DMI6000 with Hamamatsu ORCA-R2 C10600 camera and SOLA SE light source. Images were acquired in phase contrast, GFP, and Texas Red channels, and with 40x/1.30 objective. Immunostained wild-type A549, MCF-7 and HepG2 cells were imaged using Zeiss Axio Observer 7 with camera Axiocam 705 mono. Images were acquired in DIC, AF594, AF488 and DAPI channels, and with 20x/0.8 objective. The average fluorescence intensity within the nuclei of each cell in the field was calculated using the Zeiss Bio Apps Gene Expression tool. The measurement area was limited to the cell nucleus, which was detected from the signal in the DAPI-stained channel. Average fluorescence intensity in the green/red was then measured for each cell in the field and normalized to the DAPI intensity in the same cell to correct for differences in cell permeability across cell types.

### Neural network architectures

The sequence-only convolutional neural network (CNN) model aims to predict FOXA1 ChIP-seq peaks using DNA sequence input. Briefly, one-hot encoded DNA sequence input of length 240bp is first passed through a 1D convolution layer of 256 filters, where each filter is of size 24 and stride 1. After convolution, the output is processed by ReLU activation and batch normalization. A 1D max-pooling layer of size 15 and stride 15 is then applied to pool the output. The pooled output is fed into a long short-term memory (LSTM) layer to output a 32-length vector. The output vector passes through two dense layers with ReLU activation and Dropout. Finally, a single sigmoid activated linear node outputs the prediction probability.

The Bichrom models aim to assess whether chromatin features positively contribute to predicting FOXA1 binding and use a previously published interpretable bimodal neuron network architecture named Bichrom^43^. Bichrom consists of two independent sub-networks, corresponding to DNA sequence and chromatin input, respectively. The sequence sub-network is the trained CNN network described above with all the trained weights frozen. The final linear node is replaced by a new linear node activated by a tanh function. The input to the chromatin sub-network consists of the relevant chromatin feature(s) coverage track(s), binned in 20bp bins, across the same 240bp region as DNA sequence input. The chromatin feature input passes through a ReLU activated 1D convolution layer of 15 filters (kernel size 1) and a LSTM layer to output a 5-vector. A tanh activated linear node is then used to get the scalar output. The full Bichrom model works by combining the scalar values from both sub-networks into a sigmoid activated linear node to predict the TF binding label. Three Bichrom models were tested: one trained on DNA-sequence and ATAC-seq features; one trained on DNA-sequence and H3K9me3 features; and one trained on DNA-sequence and ATAC-seq, H3K9me3, H3K27ac, H3K4me1, H3K4me2, and H3K4me3 features. All chromatin features were sourced from ENCODE A549 ATAC/ChIP-seq experiments.

### Neural network training

For both model architectures, two chromosomes are held out for validation (chr11) and test (chr17). The sampling strategies differ by the model type. For the sequence-only CNN model, positive sample regions are obtained by randomly shifting 240bp long regions centered by ChIP-seq peak midpoints (-95bp<=shifting distance<95bp). Negative sample regions are sampled from four different sources: 1) flanking negative regions around ChIP-seq peaks (flanking distances: [450, -450, 500, -500, 1250, -1250, 1750, -1750]); 2) accessible regions not overlapping ChIP-seq peaks; 3) non-accessible regions not overlapping ChIP-seq peaks; 4) random regions sampled from the entire genome and not overlapping ChIP-seq peaks. The goal of sampling is to ensure the percentages of accessible regions in both positive and negative samples are the same. Bichrom models use the same positive sample regions as the sequence-only CNN model, while negative sample regions only consist of random regions sampled from the entire genome and not overlapping ChIP-seq peaks.

### Neural network feature attribution

The DeepLift-SHAP implementation from SHAP^38,39^ was employed to compute the attribution scores for each trained model. The hypothetical attribution scores were obtained by computing the DeepLift-SHAP score of all possible nucleotide choices at each base pair. Then TF-MoDISco was used to extract globally high-impact sequence patterns with the option -n 50000. The final sequence patterns were then compared to motifs from Cis-BP^89^ using Tomtom^90^.

### DNA shape analysis

The DNA shape scores were computed by DNAShapeR^91–93^ using default settings. Each type of DNA shape score was plotted around the FOXA1 motif center.

## Acknowledgements

We thank Dr. Kenneth Zaret for providing the FOXA1 bacterial expression construct, Dr. Cheng-Ming Chiang for providing the AP-1 polycistronic bacterial expression strain, and Dr. Xiaojun Lance Lian for providing the XLone piggyBac cargo strain. We also thank Dr. Yanming Wang for providing us with A549 human lung carcinoma cells. We are grateful to Dr. Cheryl Keller and others in the Huck Genomics Research Incubator for assistance, training, and discussions related to high-throughput sequencing. We acknowledge all members in the Bai lab for insightful comments on the manuscript. We also thank the members of the Center of Eukaryotic Gene Regulation at Pennsylvania State University for discussions and technical support. This work is supported by the National Institutes of Health (T32 GM125592 to H.K. and E.L., R35 GM127034 to S.T., R35 GM144135 to S.M., and R35 GM139654 to L.B.) and the Graduate Research Innovation fund from the Huck Institute of Life Sciences (to C.X.).

---

**Extended Data Fig. 1.**
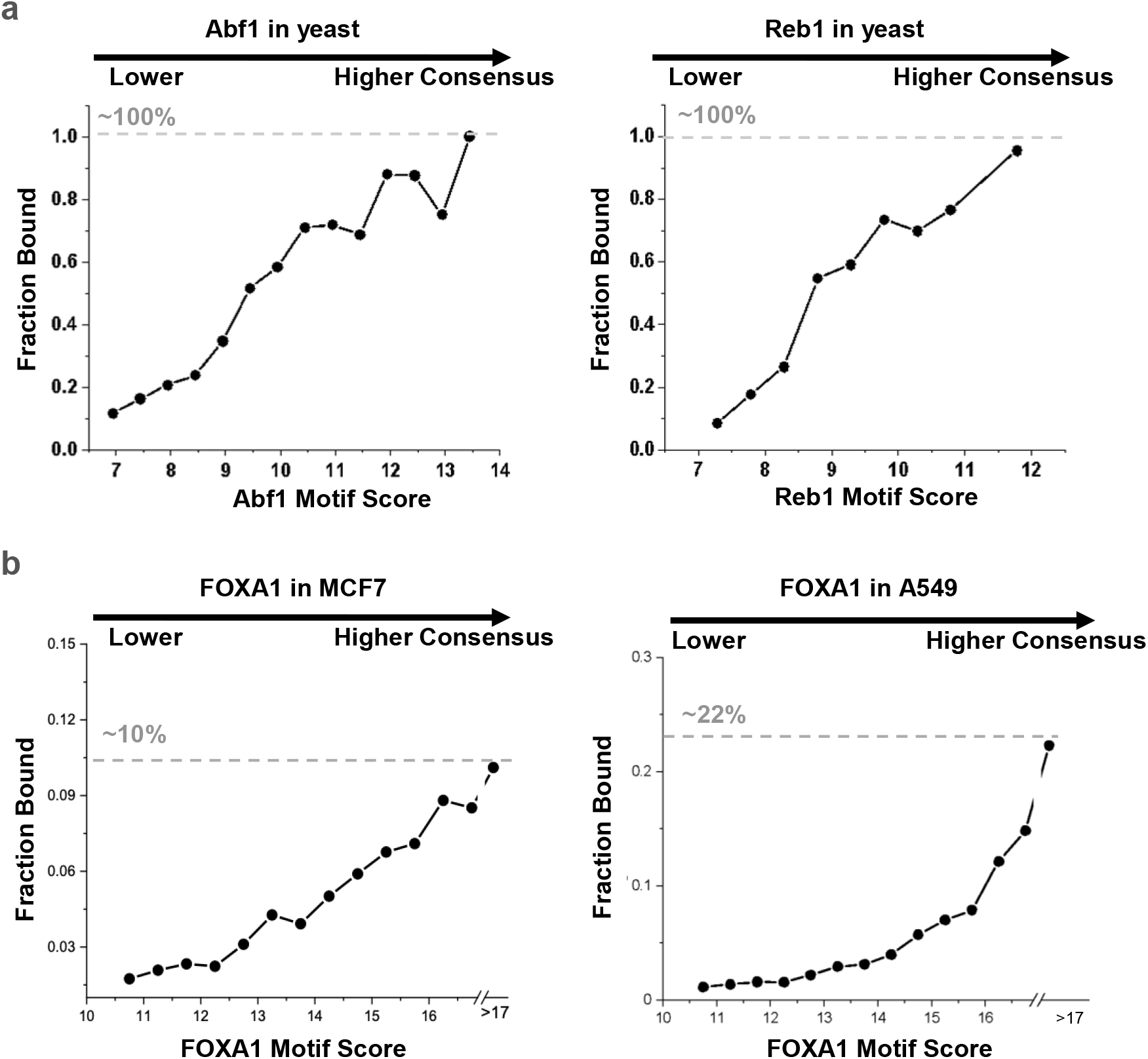
PFs in yeast, but not in mammalian cells, bind to a large fraction of their consensus. **a)** Bound fraction as a function of motif score for yeast PFs, Abf1 (left) and Reb1 (right). Dotted line represents the fraction of perfect consensus motifs that are occupied by the corresponding PFs. Abf1 and Reb1 binding data are from Ref 29. **b**, Same as panel a, but for human pioneer factor FOXA1 in MCF-7 cells (left) and A549 cells (right).

**Extended Data Fig. 2.**
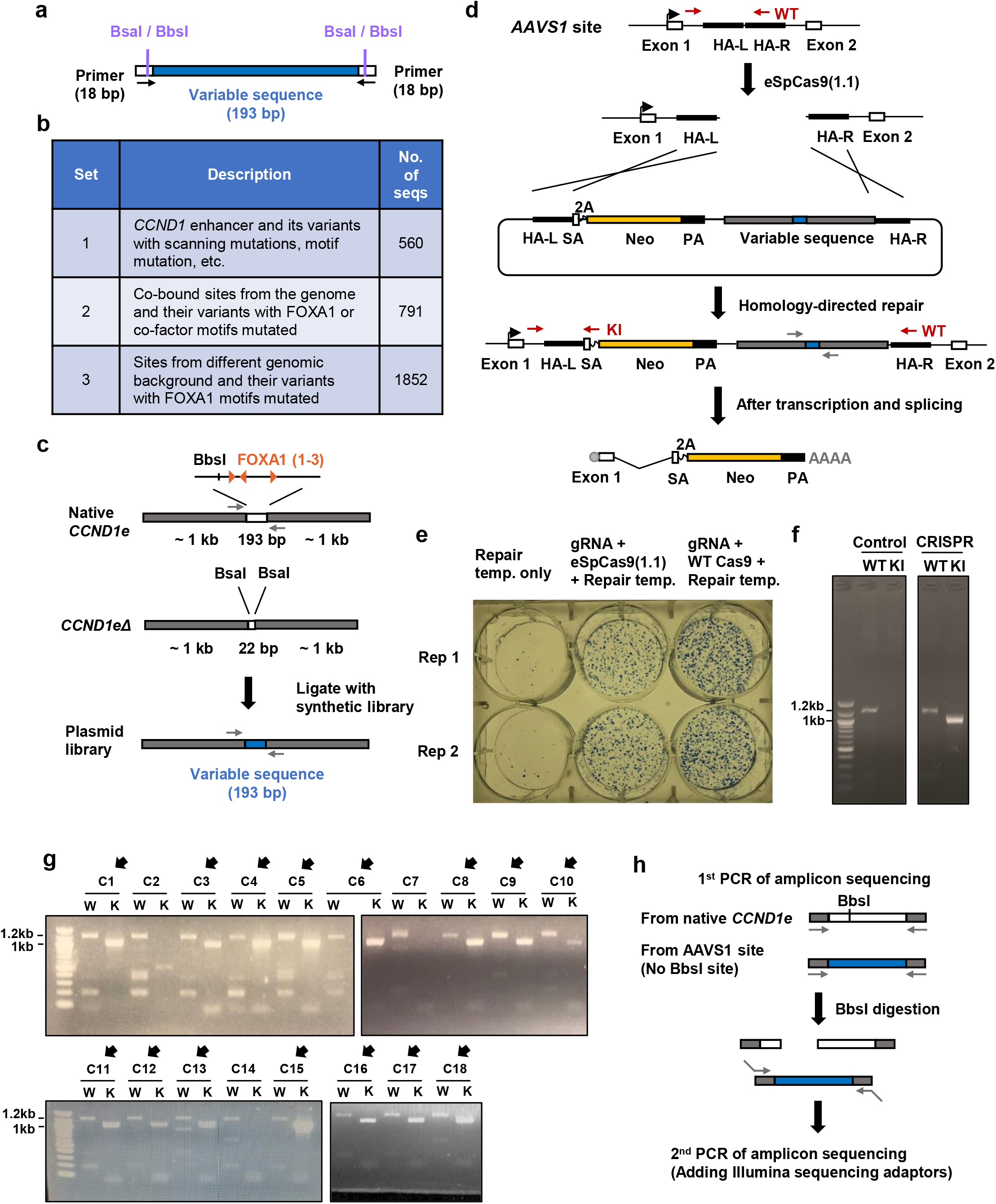
ChIP-ISO procedure and quality test. **a)** Diagram of ChIP-ISO library oligonucleotide design. The variable region of each sequence spans 193 bp, as indicated in blue. The BsaI / BbsI is the restriction enzyme recognition sites engineered in the 18 bp flanking primer sequences. **b)** Table summarizing the three subsets of our ChIP-ISO oligonucleotide library. **c)** Plasmid construction strategy. We start off with the native *CCND1* enhancer sequence, cloned into the pAAVS1-Nst-MCS plasmid background. We then delete a 193 bp region containing all three FOXA1 binding sites, as well as a BbsI cutting site, and replace it with two BsaI cutting sites. Finally, we ligate the 193 bp ChIP-ISO oligonucleotides (blue) into this plasmid using the BsaI sites. Grey arrows represent primers used for ChIP-ISO amplicon sequencing. **d)** Strategy for integration of the plasmid library into the AAVS1 site using CRISPR/Cas9. Double-strand break is generated at the AAVS1 site by pSpCas9 (1.1). The region containing library and background sequences, as well as the selection marker (Neo), is integrated into the *AAVS1* site (inside the first intron of *PPP1R12C* gene) by homology-directed repair. If integrated successfully, the promoterless Neo gene will be transcribed together with the endogenous *PPP1R12C* gene, and RNA splicing will happen between exon 1 and the splicing acceptor (SA) site. A 2A peptide is engineered before the Neo gene to make sure the translated Neo protein is folded independently. HA-L, left homologous arm; HA-R, right homologous arm. WT, a primer pair amplifying unintegrated *AAVS1* site; KI, a primer pair amplifying *AAVS1* site with successful integration. **e)** A549 colonies after integration with no Cas9 (left, control), eSpCas9, and wt Cas9. The colonies are selected with G418 for 12 days, and stained with methylene blue. **f)** PCR test on mixed colonies after integration. The “KI” primer pair (see d) amplifies across the integration junction, and therefore the band is only visible when the plasmid sequence is integrated into the right locus. **g)** Same as in panel f except that the PCR is carried out in 18 single colonies. 15 out of the 18 colonies show the right PCR band. **h)** Strategy of amplicon sequencing. After first round of PCR using the primer pair in panel c, we perform BbsI digestion, which cuts the native *CCND1* enhancers, but not the synthetic ones. The second round PCR (with sequencing adaptors) can therefore selectively amplify the ISO library for sequencing.

**Extended Data Fig. 3.**
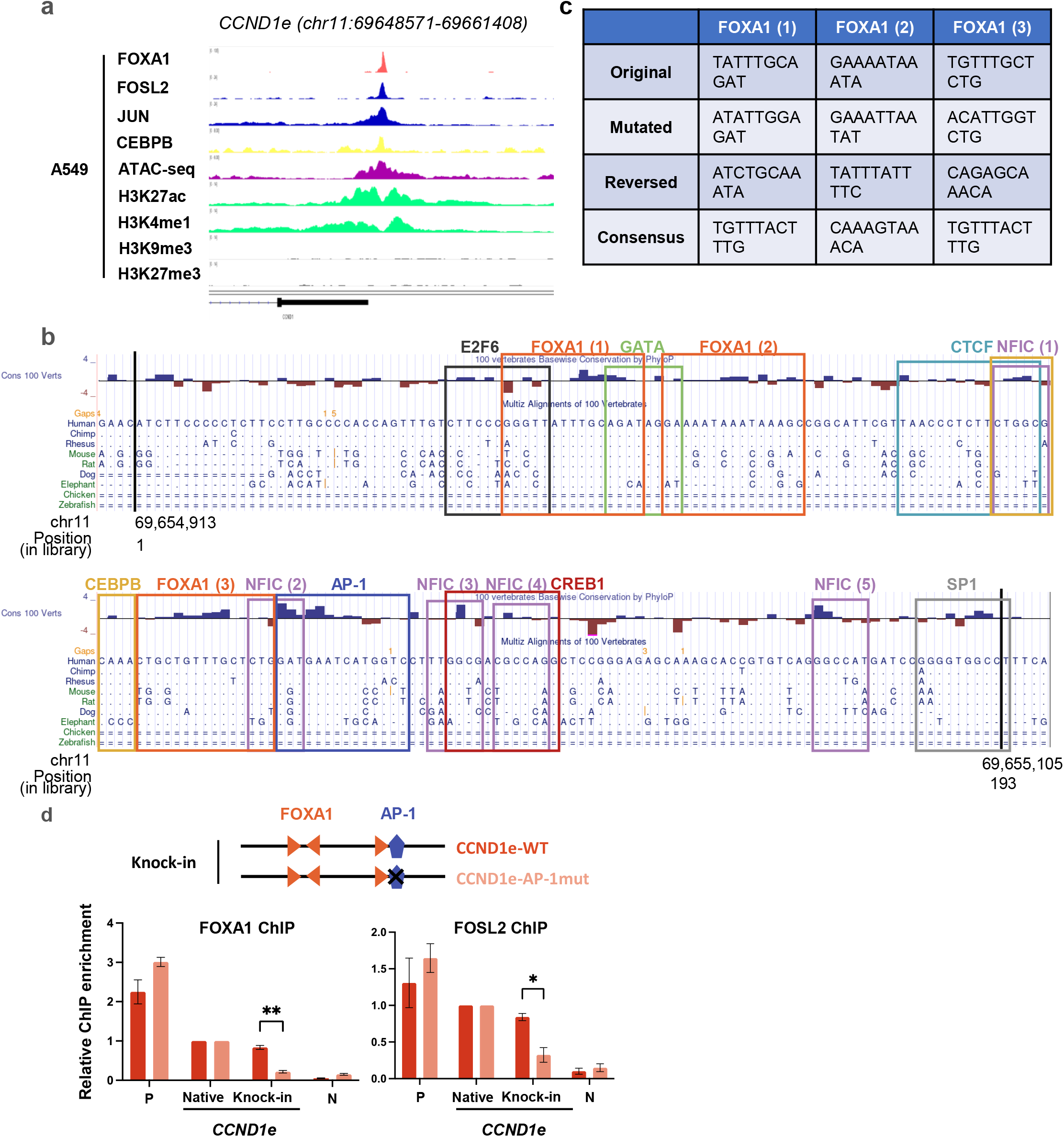
*CCND1* enhancer sequence and low-throughput test. **a)** Genomic tracks of TF and histone modification ChIP-seq and ATAC-seq signals at the *CCND1* enhancer in WT A549 cells. **b)** Conservation of DNA sequence within the ∼200 bp *CCND1* enhancer. Blue: conserved nucleotide, brown: non-conserved. TF motifs are outlined and color coded. Vertical black lines marks the exact 193 bp sequences use in our ISO library. **c)** Manipulation of the FOXA1 motifs in the *CCND1* enhancer. Each motif is mutated, orientation reversed, or converted into a perfect consensus. **d)** Low-throughput FOXA1 and FOSL2 ChIP-qPCR for three biological replicates showing the impact of the AP-1 motif on their binding. P: positive control, Native: native *CCND1* enhancer, Knock-in = AAVS1-integrated *CCND1* enhancer (wt or AP-1 mut), N: negative control. *: p < 0.05, **: p < 0.01.

**Extended Data Fig. 4.**
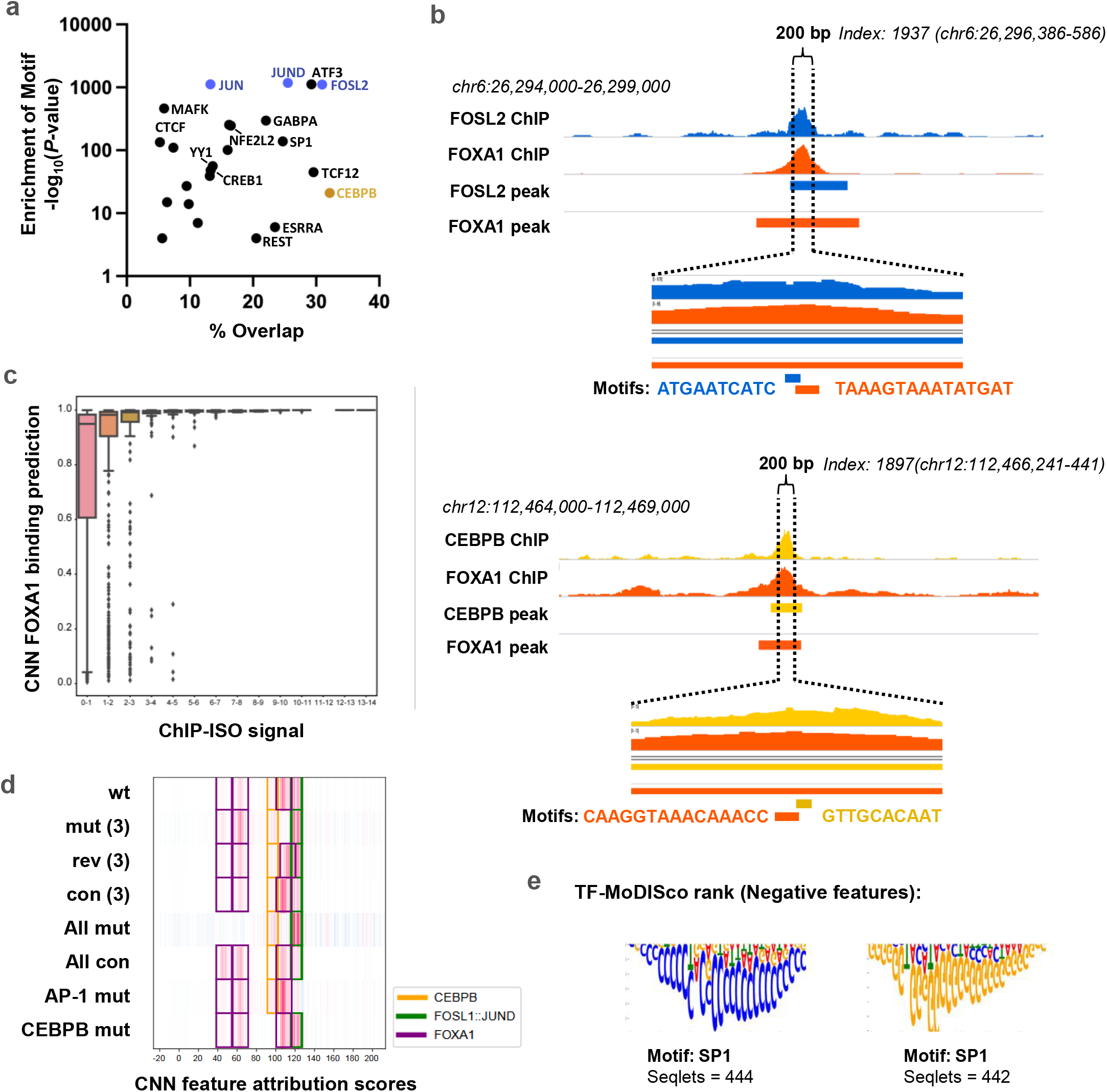
Genome-wide FOX1 / TF co-binding. **a)** The same as Fig. 3a lower panel, except the x axis represents the percentage of a TF ChIP-seq peaks overlapped with FOXA1 peaks (Fig. 3a is the percentage of the overlapped FOXA1 peaks). **b)** Two examples of native sequence with co-binding events included in our ISO library. Top: a sequence with both FOSL2 and FOXA1 ChIP-seq peaks and their motifs in proximity. Dotted lines represent the boundaries of the ∼200 bp library sequence. Blue bar: FOSL2 motif, orange bar: FOXA1 motif. Bottom: Same as top, but for a sequence with CEBPB / FOXA1 overlap. **c)** Histogram showing distributions of scores given by a sequence-trained CNN to sequences that were tested by ChIP-ISO, where the sequences are grouped according to ChIP-ISO signal. The CNN is trained to predict FOXA1 ChIP-seq data in A549 cells. **d)** Heatmap showing DeepLIFT-Shap feature attribution scores from the sequence-trained CNN at eight *CCND1* enhancer variants. **e)** Top two motifs detected by TF-MoDISco in the genome-wide DeepLIFT-Shap negative feature attribution scores. The TF family of matching motifs is annotated for each motif, as is the number of seqlets used by TF-MoDISco to construct each motif.

**Extended Data Fig. 5.**
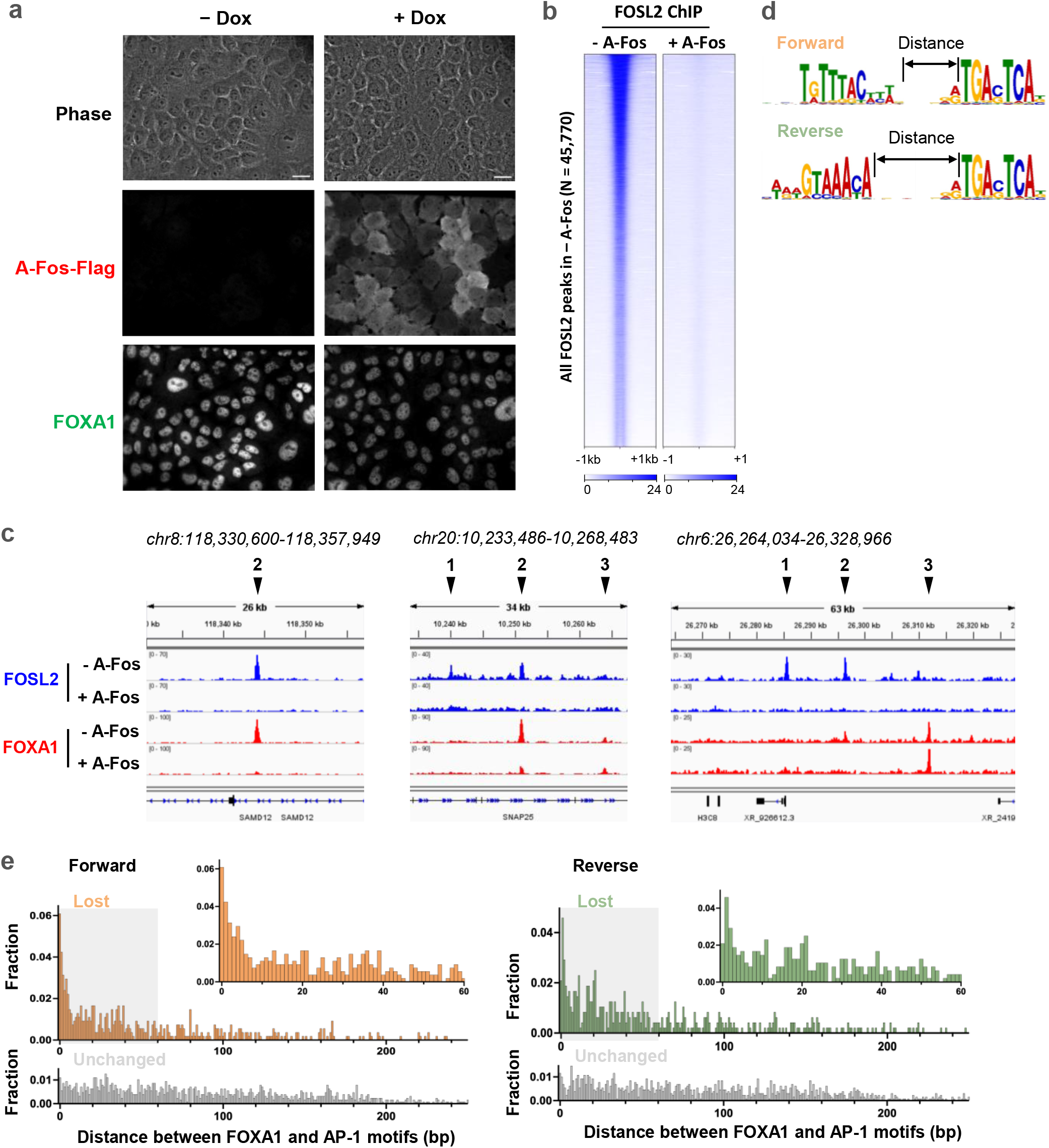
Imaging and genomic data ± A-Fos induction. **a)** Immunostaining of FLAG-tagged A-Fos and FOXA1 ± A-Fos induction. Scale bar: 20 μm. **b)** Heatmaps showing FOSL2 ChIP-seq signal across genome-wide FOSL2 binding sites ± A-Fos induction. **c)** More genomic tracks of FOSL2 and FOXA1 ChIP-seq ± A-Fos induction. Arrows 1-3 demarcate examples of AP-1 unique, AP-1 / FOXA1 overlapped, and FOXA1 unique sites, respectively. **d)** Definition of the forward and reverse orientation between FOXA1 and AP-1 motifs, as well as the distance in between. **e)** Distribution of the distances between FOXA1 and AP-1 motifs in the lost and unchanged sites upon A-Fos induction. The motif pairs with two different orientations are separately analyzed.

**Extended Data Fig. 6.**
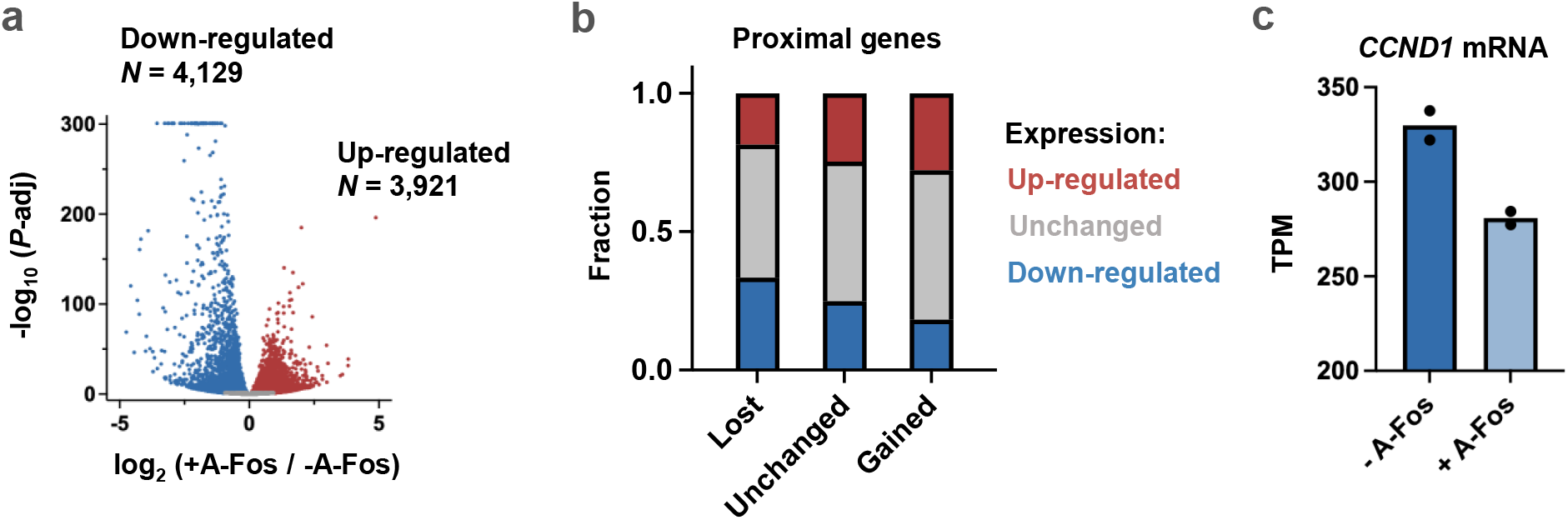
Differential expression analysis of the RNAseq ± A-Fos induction. **a)** Volcano plot of RNAseq counts in ± A-Fos conditions. Red and blue represent up-regulated vs down-regulated genes in the presence of A-Fos induction, respectively. **b)** Differential regulation of genes proximal to lost, unchanged, and gained FOXA1 peaks. The genes close to the lost peaks are more likely to be down-regulated. **c)** *CCND1* mRNA is downregulated in the presence of A-Fos, consistent with the lost of FOXA1 binding in the *CCND1* enhancer.

**Extended Data Fig. 7.**
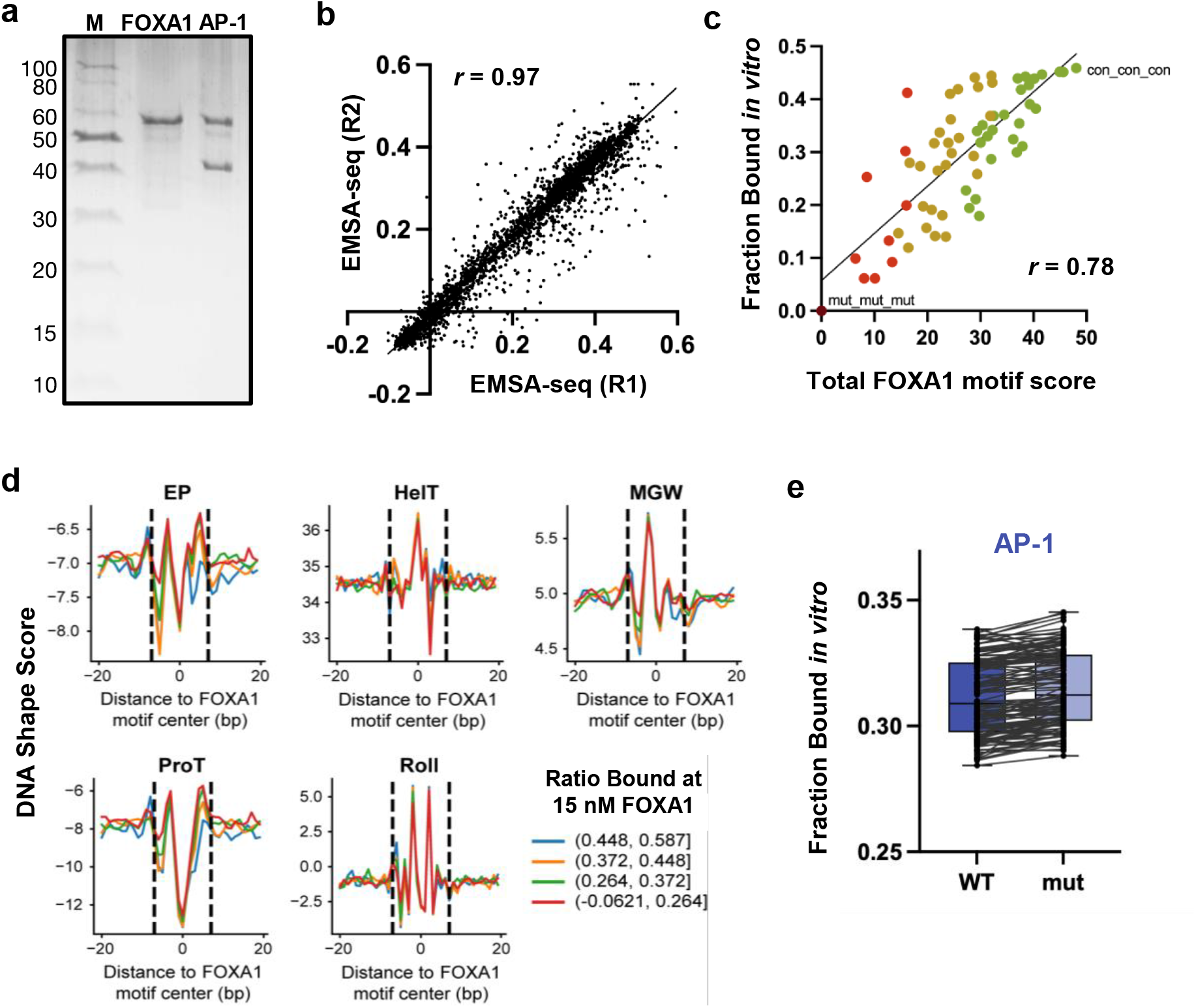
More information related to *in vitro* EMSA-seq experiment. **a)** Gel of recombinant proteins. Purified proteins were run on an 18% SDS-PAGE gel and stained with Coomassie Brilliant Blue. M: Molecular weight marker. **b)** Correlation between two EMSA-seq biological replicates. Both x and y axis shows the fraction bound at 15 nM FOXA1. **c)** Correlation between *in vitro* binding strength (fraction bound at 15 nM FOXA1) and total FOXA1 motif score in the *CCND1* enhancer subset of the ISO library. Green, yellow, orange, and red dots correspond to *CCND1* variants with three, two, one, or zero FOXA1 motifs. **d)** Plots showing five DNA shape parameters averaged across four sets of FOXA1 binding sites. DNA shape parameters include electrostatic potential (EP), helix twist (HelT), minor groove width (MGW), propeller twist (ProT), and roll. Four colors represent different range of *in vitro* binding strength. **e)** Box-and-whisker plots showing the changes of FOXA1 EMSA-seq signal on templates ±AP-1 motif.

**Extended Data Fig. 8.**
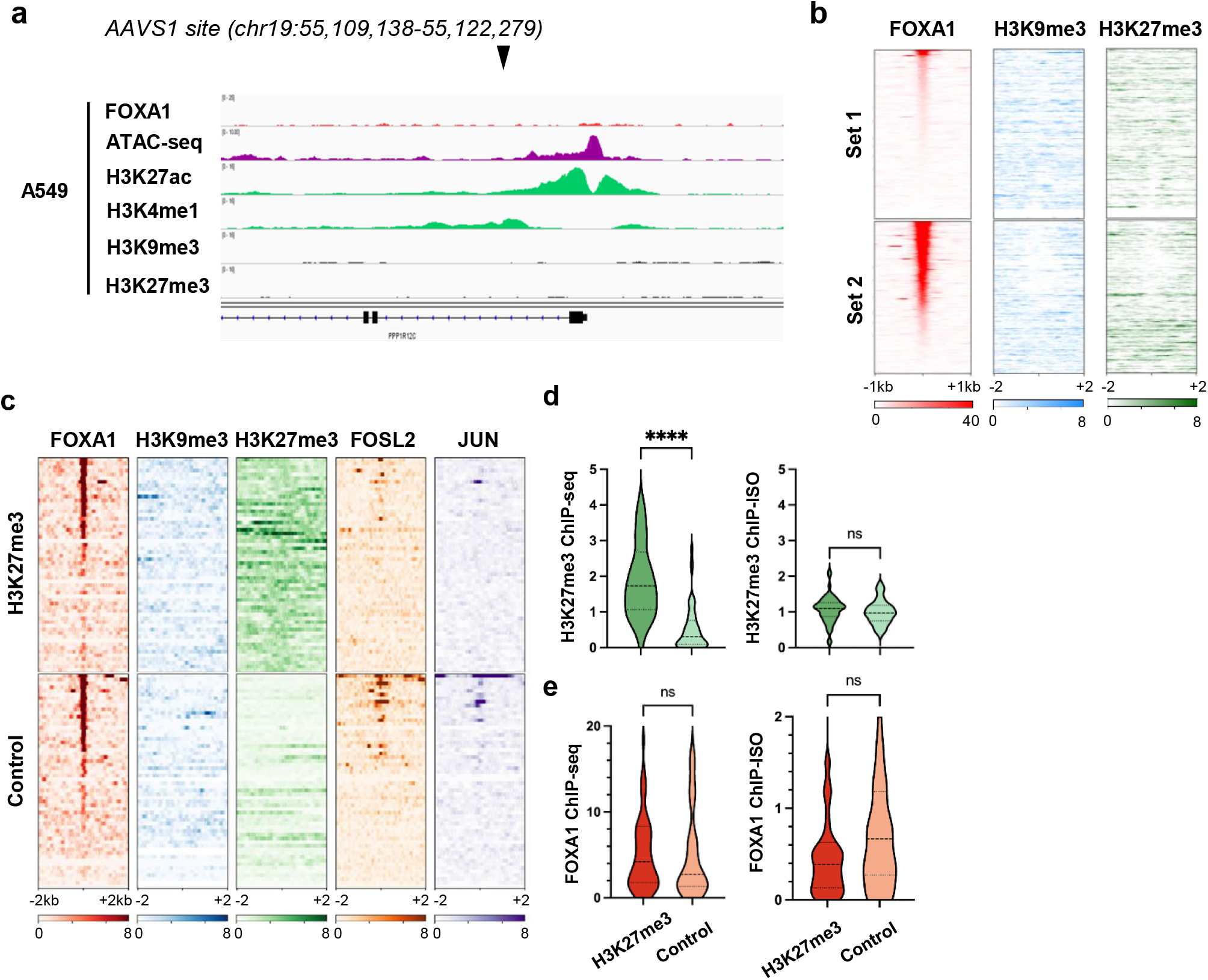
The effect of chromatin states on FOXA1 binding. **a)** Genomic tracks of TF and histone modification ChIP-seq and ATAC-seq signals at the *AAVS1* safe-harbor locus in WT A549 cells. Arrow indicates locus where library sequences were inserted. The insertion site is near active histone marks, but not the repressive ones. **b)** ChIP-seq signals of repressive histone marks near the native sequences included in our ISO library in Fig. 6A. **c)** Heatmap of FOXA1, H3K9me3, H3K27me3, FOSL2, and Jun ChIP-seq signals for a set of ChIP-ISO library sequences derived from H3K27me3-marked genomic loci (top) and a set of control sequences with comparable FOXA1 binding level derived from euchromatic loci (bottom). **d)** Violin plot of H3K27me3 ChIP-seq signals for sequences in panel c at their native genomic loci (left) and ChIP-ISO signals of the same sequences at AAVS1 (right). Dark green: H3K27me3 set, and light green: control set from panel c. ****: p < 0.0001 and ns: non-significant. **e)** Same as panel d, but for FOXA1.

**Extended Data Fig. 9.**
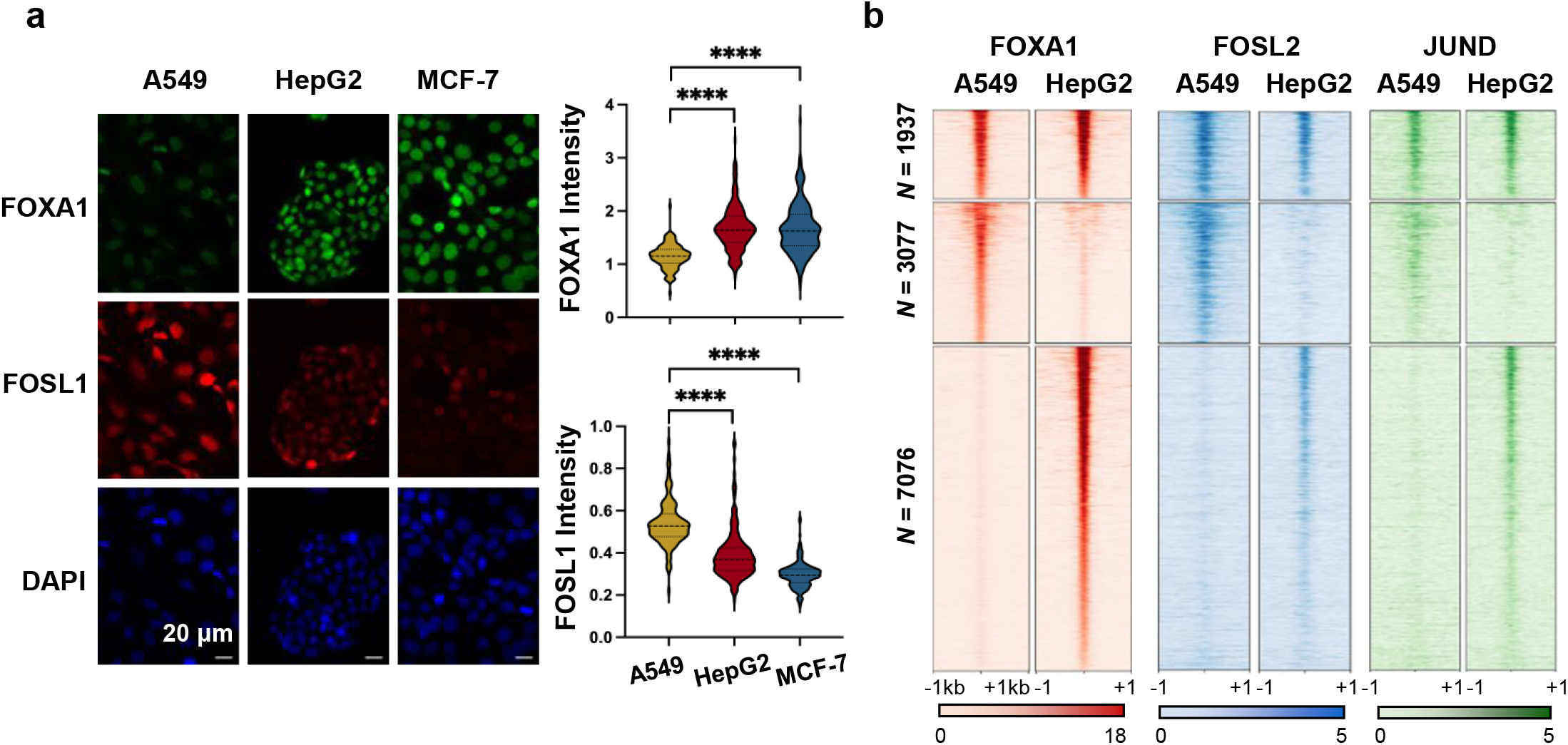
Differential binding of FOXA1 in HepG2 and MCF-7. **a)** Immunostaining of FOXA1 (green), FOSL1 (red), and nuclei (DAPI) in fixed A549, HepG2, and MCF-7 cells. Right: Violin plots showing normalized FOXA1 (top) and FOSL1 (bottom) intensities in these three cell lines. Scale bar: 20 μm. **b)** Differential FOXA1 binding analysis in A549 and HepG2 cells, with heatmaps showing common (upper), A549-specific (middle), and HepG2-specific (bottom) peaks. FOSL2 and JUND ChIP-seq signals over the same regions are shown on the right.

**Extended Data Fig. 10.**
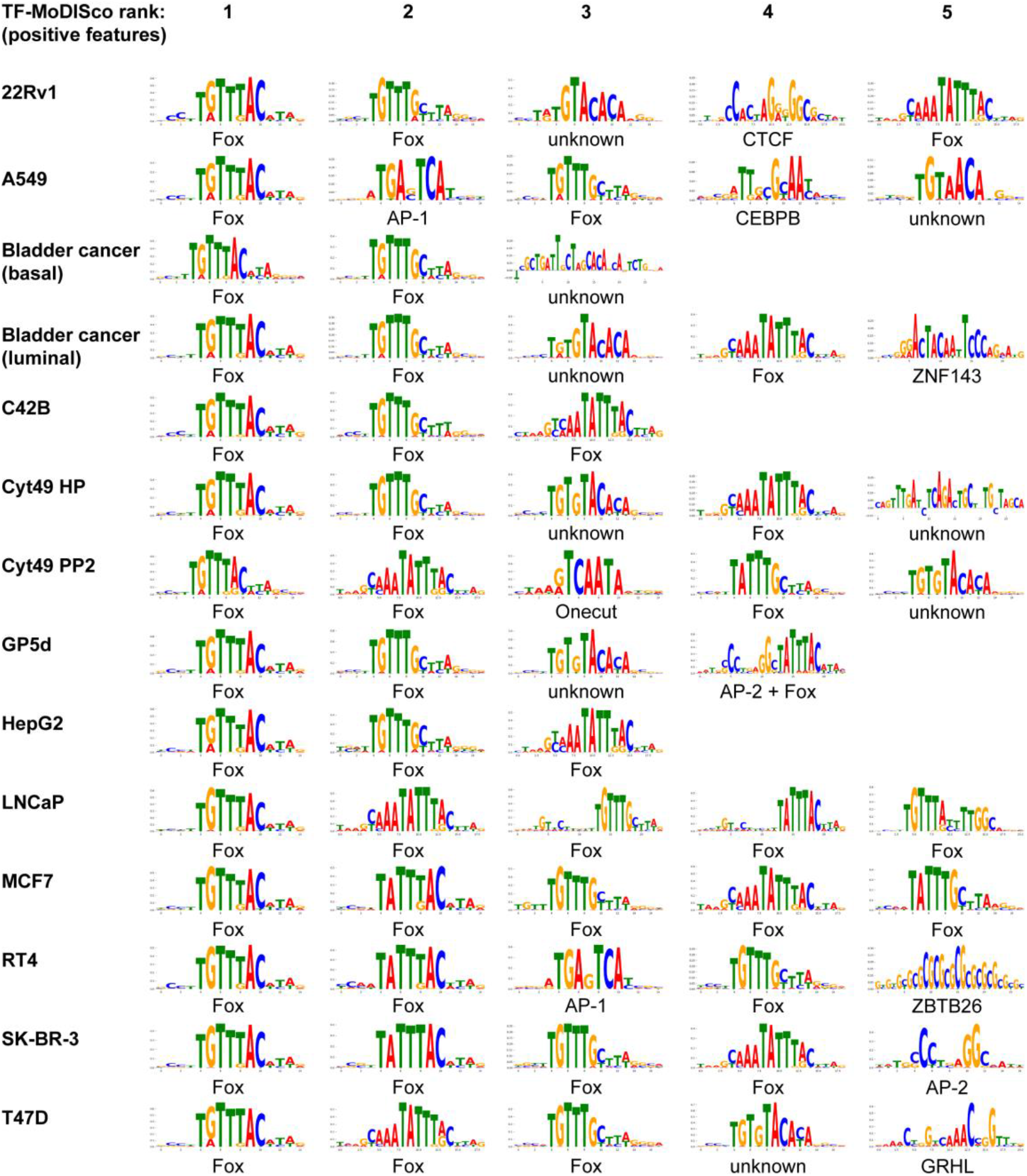
Motifs that are predictive of FOXA1 binding in sequence-trained CNNs. Top ranking motifs detected by TF-MoDISco in the genome-wide DeepLIFT-Shap positive feature attribution scores at sites predicted by a sequence-trained CNN to be bound by FOXA1 in a selection of cell types for which FOXA1 ChIP-seq data is available. The top five ranking motifs are shown, unless TF-MoDISco returned fewer than five motifs. The TF family of best-matching motifs is annotated for each motif.

## References

1. Wang, J. et al. Sequence features and chromatin structure around the genomic regions bound by 119 human transcription factors. Genome Res 22, 1798–812 (2012).

2. Arvey, A., Agius, P., Noble, W.S. & Leslie, C. Sequence and chromatin determinants of cell-type-specific transcription factor binding. Genome Res 22, 1723–34 (2012).

3. Spitz, F. & Furlong, E.E. Transcription factors: from enhancer binding to developmental control. Nat Rev Genet 13, 613–26 (2012).

4. Gordan, R. et al. Genomic regions flanking E-box binding sites influence DNA binding specificity of bHLH transcription factors through DNA shape. Cell Rep 3, 1093–104 (2013).

5. Abe, N. et al. Deconvolving the recognition of DNA shape from sequence. Cell 161, 307–18 (2015).

6. Jolma, A. et al. DNA-dependent formation of transcription factor pairs alters their binding specificity. Nature 527, 384–8 (2015).

7. Donaghey, J. et al. Genetic determinants and epigenetic effects of pioneer-factor occupancy. Nat Genet 50, 250–258 (2018).

8. Kim, S. et al. DNA-guided transcription factor cooperativity shapes face and limb mesenchyme. bioRxiv (2023).

9. Domcke, S. et al. Competition between DNA methylation and transcription factors determines binding of NRF1. Nature 528, 575–9 (2015).

10. Kaluscha, S. et al. Evidence that direct inhibition of transcription factor binding is the prevailing mode of gene and repeat repression by DNA methylation. Nat Genet 54, 1895–1906 (2022).

11. Barozzi, I. et al. Coregulation of transcription factor binding and nucleosome occupancy through DNA features of mammalian enhancers. Mol Cell 54, 844–857 (2014).

12. Neikes, H.K. et al. Quantification of absolute transcription factor binding affinities in the native chromatin context using BANC-seq. Nat Biotechnol (2023).

13. Soufi, A., Donahue, G. & Zaret, K.S. Facilitators and impediments of the pluripotency reprogramming factors’ initial engagement with the genome. Cell 151, 994–1004 (2012).

14. Sinha, K.K., Bilokapic, S., Du, Y., Malik, D. & Halic, M. Histone modifications regulate pioneer transcription factor cooperativity. Nature 619, 378–384 (2023).

15. Kim, S. & Shendure, J. Mechanisms of Interplay between Transcription Factors and the 3D Genome. Mol Cell 76, 306–319 (2019).

16. Garcia, D.A. et al. An intrinsically disordered region-mediated confinement state contributes to the dynamics and function of transcription factors. Mol Cell 81, 1484–1498 e6 (2021).

17. Brodsky, S., Jana, T. & Barkai, N. Order through disorder: The role of intrinsically disordered regions in transcription factor binding specificity. Curr Opin Struct Biol 71, 110–115 (2021).

18. Slattery, M. et al. Absence of a simple code: how transcription factors read the genome. Trends Biochem Sci 39, 381–99 (2014).

19. Keilwagen, J., Posch, S. & Grau, J. Accurate prediction of cell type-specific transcription factor binding. Genome Biol 20, 9 (2019).

20. Luo, Y., North, J.A., Rose, S.D. & Poirier, M.G. Nucleosomes accelerate transcription factor dissociation. Nucleic Acids Res 42, 3017–27 (2014).

21. Soufi, A. et al. Pioneer transcription factors target partial DNA motifs on nucleosomes to initiate reprogramming. Cell 161, 555–568 (2015).

22. Donovan, B.T. et al. Basic helix-loop-helix pioneer factors interact with the histone octamer to invade nucleosomes and generate nucleosome-depleted regions. Mol Cell 83, 1251–1263 e6 (2023).

23. Guan, R., Lian, T., Zhou, B.R., Wheeler, D. & Bai, Y. Structural mechanism of LIN28B nucleosome targeting by OCT4. Mol Cell 83, 1970–1982 e6 (2023).

24. Donovan, B.T., Chen, H., Jipa, C., Bai, L. & Poirier, M.G. Dissociation rate compensation mechanism for budding yeast pioneer transcription factors. Elife 8(2019).

25. Zaret, K.S. & Carroll, J.S. Pioneer transcription factors: establishing competence for gene expression. Genes Dev 25, 2227–41 (2011).

26. Balsalobre, A. & Drouin, J. Pioneer factors as master regulators of the epigenome and cell fate. Nat Rev Mol Cell Biol 23, 449–464 (2022).

27. Bulyk, M.L., Drouin, J., Harrison, M.M., Taipale, J. & Zaret, K.S. Pioneer factors - key regulators of chromatin and gene expression. Nat Rev Genet (2023).

28. Yan, C., Chen, H. & Bai, L. Systematic Study of Nucleosome-Displacing Factors in Budding Yeast. Mol Cell 71, 294–305 e4 (2018).

29. Rossi, M.J. et al. A high-resolution protein architecture of the budding yeast genome. Nature 592, 309–314 (2021).

30. Bernardo, G.M. & Keri, R.A. FOXA1: a transcription factor with parallel functions in development and cancer. Biosci Rep 32, 113–30 (2012).

31. Cirillo, L.A. et al. Opening of compacted chromatin by early developmental transcription factors HNF3 (FoxA) and GATA-4. Mol Cell 9, 279–89 (2002).

32. Fakhouri, T.H., Stevenson, J., Chisholm, A.D. & Mango, S.E. Dynamic chromatin organization during foregut development mediated by the organ selector gene PHA-4/FoxA. PLoS Genet 6(2010).

33. Serandour, A.A. et al. Epigenetic switch involved in activation of pioneer factor FOXA1-dependent enhancers. Genome Res 21, 555–65 (2011).

34. Lupien, M. et al. FoxA1 translates epigenetic signatures into enhancer-driven lineage-specific transcription. Cell 132, 958–70 (2008).

35. Wang, H. et al. A systematic approach identifies FOXA1 as a key factor in the loss of epithelial traits during the epithelial-to-mesenchymal transition in lung cancer. BMC Genomics 14, 680 (2013).

36. Li, J., Zhang, S., Zhu, L. & Ma, S. Role of transcription factor FOXA1 in non-small cell lung cancer. Mol Med Rep 17, 509–521 (2018).

37. Eeckhoute, J., Carroll, J.S., Geistlinger, T.R., Torres-Arzayus, M.I. & Brown, M. A cell-type-specific transcriptional network required for estrogen regulation of cyclin D1 and cell cycle progression in breast cancer. Genes Dev 20, 2513–26 (2006).

38. Shrikumar, A., Greenside, P. & Kundaje, A. Learning Important Features Through Propagating Activation Differences. International Conference on Machine Learning, *Vol* 70 70(2017).

39. Lundberg, S.M. & Lee, S.I. A Unified Approach to Interpreting Model Predictions. Advances in Neural Information Processing Systems 30 (Nips 2017) 30(2017).

40. Shrikumar, A. et al. Technical note on transcription factor motif discovery from importance scores (TF-MoDISco) version 0.5. 6.5. arXiv preprint arXiv:1811.00416 (2018).

41. Olive, M. et al. A dominant negative to activation protein-1 (AP1) that abolishes DNA binding and inhibits oncogenesis. J Biol Chem 272, 18586–94 (1997).

42. Biddie, S.C. et al. Transcription factor AP1 potentiates chromatin accessibility and glucocorticoid receptor binding. Mol Cell 43, 145–55 (2011).

43. Srivastava, D., Aydin, B., Mazzoni, E.O. & Mahony, S. An interpretable bimodal neural network characterizes the sequence and preexisting chromatin predictors of induced transcription factor binding. Genome Biol 22, 20 (2021).

44. Kleinschmidt, H., Xu, C. & Bai, L. Using Synthetic DNA Libraries to Investigate Chromatin and Gene Regulation. Chromosoma 132, 167–189 (2023).

45. Teytelman, L., Thurtle, D.M., Rine, J. & van Oudenaarden, A. Highly expressed loci are vulnerable to misleading ChIP localization of multiple unrelated proteins. Proc Natl Acad Sci U S A 110, 18602–7 (2013).

46. Swinstead, E.E. et al. Steroid Receptors Reprogram FoxA1 Occupancy through Dynamic Chromatin Transitions. Cell 165, 593–605 (2016).

47. Geusz, R.J. et al. Sequence logic at enhancers governs a dual mechanism of endodermal organ fate induction by FOXA pioneer factors. Nat Commun 12, 6636 (2021).

48. Farley, E.K. et al. Suboptimization of developmental enhancers. Science 350, 325–8 (2015).

49. Crocker, J. et al. Low affinity binding site clusters confer hox specificity and regulatory robustness. Cell 160, 191–203 (2015).

50. Hovland, A.S. et al. Pluripotency factors are repurposed to shape the epigenomic landscape of neural crest cells. Dev Cell 57, 2257–2272 e5 (2022).

51. Bejjani, F., Evanno, E., Zibara, K., Piechaczyk, M. & Jariel-Encontre, I. The AP-1 transcriptional complex: Local switch or remote command? Biochim Biophys Acta Rev Cancer 1872, 11–23 (2019).

52. Fu, X. et al. FOXA1 upregulation promotes enhancer and transcriptional reprogramming in endocrine-resistant breast cancer. Proc Natl Acad Sci U S A 116, 26823–26834 (2019).

53. Bi, M. et al. Enhancer reprogramming driven by high-order assemblies of transcription factors promotes phenotypic plasticity and breast cancer endocrine resistance. Nat Cell Biol 22, 701–715 (2020).

54. Milan, M. et al. FOXA2 controls the cis-regulatory networks of pancreatic cancer cells in a differentiation grade-specific manner. EMBO J 38, e102161 (2019).

55. Wolf, B.K. et al. Cooperation of chromatin remodeling SWI/SNF complex and pioneer factor AP-1 shapes 3D enhancer landscapes. Nat Struct Mol Biol 30, 10–21 (2023).

56. Vierbuchen, T. et al. AP-1 Transcription Factors and the BAF Complex Mediate Signal-Dependent Enhancer Selection. Mol Cell 68, 1067–1082 e12 (2017).

57. Avsec, Z. et al. Base-resolution models of transcription-factor binding reveal soft motif syntax. Nat Genet 53, 354–366 (2021).

58. de Almeida, B.P., Reiter, F., Pagani, M. & Stark, A. DeepSTARR predicts enhancer activity from DNA sequence and enables the de novo design of synthetic enhancers. Nat Genet 54, 613–624 (2022).

59. Mirny, L.A. Nucleosome-mediated cooperativity between transcription factors. Proc Natl Acad Sci U S A 107, 22534–9 (2010).

60. Mayran, A. et al. Pioneer factor Pax7 deploys a stable enhancer repertoire for specification of cell fate. Nat Genet 50, 259–269 (2018).

61. Zaret, K.S. Pioneer Transcription Factors Initiating Gene Network Changes. Annu Rev Genet 54, 367–385 (2020).

62. Whitton, H. et al. Changes at the nuclear lamina alter binding of pioneer factor Foxa2 in aged liver. Aging Cell 17, e12742 (2018).

63. Chen, H., Yan, C., Dhasarathy, A., Kladde, M. & Bai, L. Investigating pioneer factor activity and its coordination with chromatin remodelers using integrated synthetic oligo assay. STAR Protoc 4, 102279 (2023).

64. Grant, C.E., Bailey, T.L. & Noble, W.S. FIMO: scanning for occurrences of a given motif. Bioinformatics 27, 1017–8 (2011).

65. Quinlan, A.R. & Hall, I.M. BEDTools: a flexible suite of utilities for comparing genomic features. Bioinformatics 26, 841–2 (2010).

66. McLeay, R.C. & Bailey, T.L. Motif Enrichment Analysis: a unified framework and an evaluation on ChIP data. BMC Bioinformatics 11, 165 (2010).

67. Ramirez, F. et al. deepTools2: a next generation web server for deep-sequencing data analysis. Nucleic Acids Res 44, W160–5 (2016).

68. Randolph, L.N., Bao, X., Zhou, C. & Lian, X. An all-in-one, Tet-On 3G inducible PiggyBac system for human pluripotent stem cells and derivatives. Sci Rep 7, 1549 (2017).

69. Hazelbaker, D.Z. et al. A multiplexed gRNA piggyBac transposon system facilitates efficient induction of CRISPRi and CRISPRa in human pluripotent stem cells. Sci Rep 10, 635 (2020).

70. Chen, S., Zhou, Y., Chen, Y. & Gu, J. fastp: an ultra-fast all-in-one FASTQ preprocessor. Bioinformatics 34, i884–i890 (2018).

71. Martin, M. Cutadapt removes adapter sequences from high-throughput sequencing reads. *EMBnet*. journal 17, 10–12 (2011).

72. Gaspar, J.M. NGmerge: merging paired-end reads via novel empirically-derived models of sequencing errors. BMC Bioinformatics 19, 536 (2018).

73. Vasimuddin, M., Misra, S., Li, H. & Aluru, S. Efficient Architecture-Aware Acceleration of BWA-MEM for Multicore Systems. 2019 Ieee 33rd International Parallel and Distributed Processing Symposium (Ipdps 2019), 314–324 (2019).

74. Barnett, D.W., Garrison, E.K., Quinlan, A.R., Stromberg, M.P. & Marth, G.T. BamTools: a C++ API and toolkit for analyzing and managing BAM files. Bioinformatics 27, 1691–2 (2011).

75. Li, H. et al. The Sequence Alignment/Map format and SAMtools. Bioinformatics 25, 2078–9 (2009).

76. Amemiya, H.M., Kundaje, A. & Boyle, A.P. The ENCODE Blacklist: Identification of Problematic Regions of the Genome. Sci Rep 9, 9354 (2019).

77. Zhang, Y. et al. Model-based analysis of ChIP-Seq (MACS). Genome Biol 9, R137 (2008).

78. Batut, B., van den Beek, M., Doyle, M.A. & Soranzo, N. RNA-Seq Data Analysis in Galaxy. Methods Mol Biol 2284, 367–392 (2021).

79. Ross-Innes, C.S. et al. Differential oestrogen receptor binding is associated with clinical outcome in breast cancer. Nature 481, 389–93 (2012).

80. O’Connor, T., Grant, C.E., Boden, M. & Bailey, T.L. T-Gene: improved target gene prediction. Bioinformatics 36, 3902–3904 (2020).

81. Zhou, Y. et al. Metascape provides a biologist-oriented resource for the analysis of systems-level datasets. Nat Commun 10 1523 (2019).

82. Fernandez Garcia, M., et al. Structural Features of Transcription Factors Associating with Nucleosome Binding. Mol Cell 75, 921–932 e6 (2019).

83. Tan, S. A modular polycistronic expression system for overexpressing protein complexes in Escherichia coli. Protein Expr Purif 21, 224–34 (2001).

84. Tan, S., Kern, R.C. & Selleck, W. The pST44 polycistronic expression system for producing protein complexes in Escherichia coli. Protein Expr Purif 40, 385–95 (2005).

85. Wang, W.M., Lee, A.Y. & Chiang, C.M. One-step affinity tag purification of full-length recombinant human AP-1 complexes from bacterial inclusion bodies using a polycistronic expression system. Protein Expr Purif 59, 144–52 (2008).

86. Ferguson, H.A. & Goodrich, J.A. Expression and purification of recombinant human c-Fos/c-Jun that is highly active in DNA binding and transcriptional activation. Nucleic Acids Research 29, art. no.-e98 (2001).

87. Robinson, M.D., McCarthy, D.J. & Smyth, G.K. edgeR: a Bioconductor package for differential expression analysis of digital gene expression data. Bioinformatics 26, 139–40 (2010).

88. Yoney, A., Bai, L., Brivanlou, A.H. & Siggia, E.D. Mechanisms underlying WNT-mediated priming of human embryonic stem cells. Development 149(2022).

89. Weirauch, M.T. et al. Determination and inference of eukaryotic transcription factor sequence specificity. Cell 158, 1431–1443 (2014).

90. Gupta, S., Stamatoyannopoulos, J.A., Bailey, T.L. & Noble, W.S. Quantifying similarity between motifs. Genome Biol 8, R24 (2007).

91. Chiu, T.P. et al. DNAshapeR: an R/Bioconductor package for DNA shape prediction and feature encoding. Bioinformatics 32, 1211–3 (2016).

92. Li, J. et al. Expanding the repertoire of DNA shape features for genome-scale studies of transcription factor binding. Nucleic Acids Res 45, 12877–12887 (2017).

93. Chiu, T.P., Rao, S., Mann, R.S., Honig, B. & Rohs, R. Genome-wide prediction of minor-groove electrostatic potential enables biophysical modeling of protein-DNA binding. Nucleic Acids Res 45, 12565–12576 (2017).

